# Magnetoelectric Nanodiscs Enable Wireless Transgene-Free Neuromodulation

**DOI:** 10.1101/2023.12.24.573272

**Authors:** Ye Ji Kim, Nicolette Driscoll, Noah Kent, Emmanuel Vargas Paniagua, Anthony Tabet, Florian Koehler, Marie Manthey, Atharva Sahasrabudhe, Lorenzo Signorelli, Danijela Gregureć, Polina Anikeeva

## Abstract

Deep-brain stimulation (DBS) with implanted electrodes revolutionized treatment of movement disorders and empowered neuroscience studies. Identifying less invasive alternatives to DBS may further extend its clinical and research applications. Nanomaterial-mediated transduction of magnetic fields into electric potentials offers an alternative to invasive DBS. Here, we synthesize magnetoelectric nanodiscs (MENDs) with a core-double shell Fe_3_O_4_-CoFe_2_O_4_-BaTiO_3_ architecture with efficient magnetoelectric coupling. We find robust responses to magnetic field stimulation in neurons decorated with MENDs at a density of 1 µg/mm^2^ despite individual-particle potentials below the neuronal excitation threshold. We propose a model for repetitive subthreshold depolarization, which combined with cable theory, corroborates our findings in vitro and informs magnetoelectric stimulation in vivo. MENDs injected into the ventral tegmental area of genetically intact mice at concentrations of 1 mg/mL enable remote control of reward behavior, setting the stage for mechanistic optimization of magnetoelectric neuromodulation and inspiring its future applications in fundamental and translational neuroscience.

## Introduction

Deep brain stimulation (DBS) via surgically implanted electrodes is a powerful therapeutic tool for neurological and psychiatric conditions^1,2^. However, the invasiveness of DBS often makes it a last-resort therapy^1,2^ and motivates vigorous investigation of less-invasive brain stimulation alternatives. Non-invasive neuromodulation approaches, such as transcranial magnetic stimulation^3–6^, temporal interference electrical stimulation^7,8^, and focused ultrasound,^9–11^ eliminate the need for implantable hardware but offer limited resolution especially in deeper brain structures, and their mechanisms of action remain a topic of investigation. Methods combining biologically imperceptible optical, magnetic, or acoustic signals with a genetic or synthetic transducer decouple penetration depth from the resolution serving as a middle ground between invasive DBS and non-invasive low-resolution alternatives. Advances in sensitive opsins recently enabled optogenetic neuromodulation with external light sources.^12,13^ However significant scattering at tissue interfaces still precludes the deployment of these tools in the deep brain, especially in larger organisms. Weak magnetic fields (MFs) take advantage of the low conductivity and magnetic permeability of biological matter to deliver signals to deep brain structures^14–16^. MFs have been transduced into mechanical torque^17,18^, heat^19–22^, and chemical release^23,24^ enabling modulation of cells expressing mechano-, thermo-, and chemo-receptors, respectively. Although genetic sensitization of identifiable cells to specified stimuli empowers fundamental neuroscience research, the need for transgenes impedes the implementation of these methods in translational studies and in clinic.

To eliminate the need for transgenes and match the effects of clinical DBS, acoustic fields and MFs have been converted directly into electrical signals via synthetic piezoelectric^25,26^ and magnetoelectric (ME)^27–29^ transducers, respectively. As compared to acoustic signals MFs offer greater penetration depth, and ME transduction at the millimeter scale has been leveraged to power miniature electrical stimulators with the added benefit of multiplexing^29–31^. To permit ME neuromodulation with transducers that can be delivered via an injection rather than a surgical placement, MF must be converted into electrical polarization at the nanoscale. However, achieving stimulation effects comparable to clinical DBS with ME nanomaterials is challenging due to their low generated potentials. Pioneering studies of ME nanoparticles (MENPs) comprising biochemically inert magnetostrictive CoFe_2_O_4_ cores and piezoelectric BaTiO_3_ shells demonstrated single-particle voltages of ∼8 µV, with the theoretical limit of ∼1 mV in physiologically safe MF conditions^28,32^. These values are 2-3 orders of magnitude lower than the changes in membrane voltage necessary to trigger action potentials in neurons^33^. Consequently, MENPs have been employed at high concentrations forming microscale aggregates with variable properties^28^.

Here, we hypothesized that enhancing potential generated by individual ME nanoparticles would enable robust, minimally invasive DBS at low concentrations of synthetic materials. To test this hypothesis, we synthesized core-double shell Fe_3_O_4_-CoFe_2_O_4_-BaTiO_3_ hexagonal nanodiscs (ME nanodiscs, MENDs), where shape anisotropy and interfacial strain in the Fe_3_O_4_-CoFe_2_O_4_ magnetostrictive interior yielded gains in strain-mediated ME coupling with the BaTiO_3_ shell and induced robust neuronal responses to MF stimuli (**Fig. 1a**). To investigate the biophysical mechanisms underlying electrical neuromodulation mediated by MENDs, we modeled neuronal excitation in the presence of weak alternating potentials. Through these mechanistic studies, we derived design rules for ME neuromodulation in vivo and demonstrated wireless control of reward behavior in mice without transgenes at MEND concentrations 100 times lower than the nanoparticle concentrations used in prior ME neuromodulation studies. Our findings pave the way for mechanism-driven optimization of ME deep-brain stimulation as tool for fundamental and preclinical neuroscience and position this approach as a minimally invasive alternative for DBS.

**Fig. 1.**
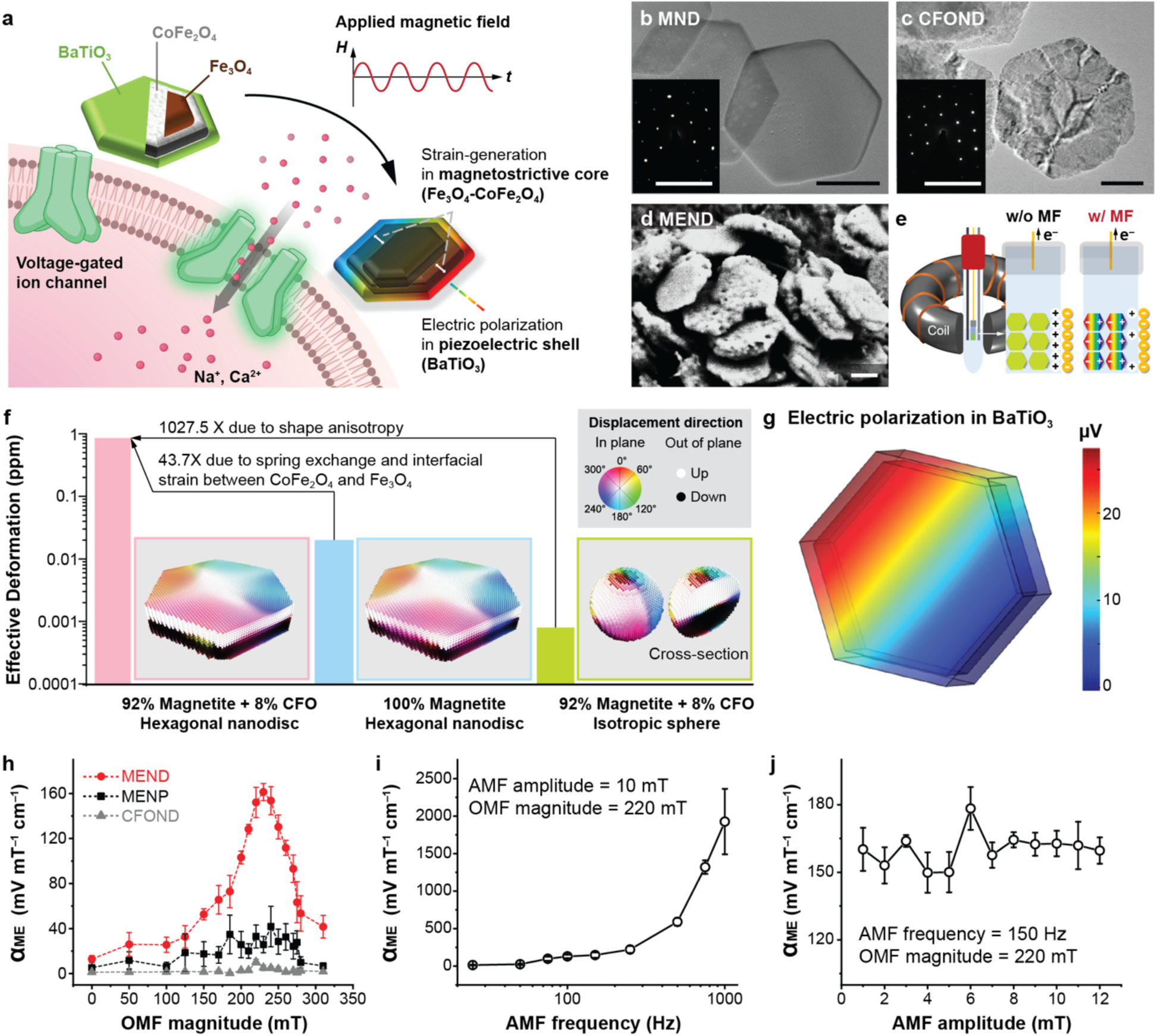
| Magnetoelectric nanodiscs for neuromodulation. a, An illustration of neuromodulation mediated by magnetoelectric nanodiscs (MEND). **b,c**, Transmission electron microscopy (TEM) images of **b,** Fe_3_O_4_ magnetic nanodiscs (MNDs), which form core of MEND and **c,** core-shell Fe_3_O_4_-CoFe_2_O_4_ nanodiscs (CFONDs). Scale bars are 100 nm. Insets show selected area electron diffraction (SAED) patterns revealing epitaxial growth. Scale bars are 10 nm^−1^. **d,** Scanning electron microscopy (SEM) image of core-double shell Fe_3_O_4_-CoFe_2_O_4_-BaTiO_3_ MENDs. Scale bar = 100 nm. **e,** Illustration of the electrochemical measurement apparatus that employs surface charge variation of MENDs in response to applied magnetic field (MF) to determine the magnetoelectric coefficient ( α*_ME_*). **f,** Simulated magnetostriction constant (bars) and magnetostrictive displacement maps (framed insets) for hexagonal CFONDs (pink), hexagonal MNDs (blue), and Fe_3_O_4_-CoFe_2_O_4_ core-shell spherical nanoparticles (green). The volume of the particles was fixed across conditions. Inset: Color index for direction in magnetostrictive displacement maps. **g,** Electric polarization generated in BaTiO_3_ shells deposited onto CFONDs upon exposure to an offset magnetic field (OMF) of 220 mT and an alternating magnetic field (AMF) with an amplitude of 10 mT. **h,** α*_ME_* at an AMF with a frequency ƒ_AMF_ =150 Hz and amplitude *H*_AMF_ = 10 mT measured at varying magnitudes of OMF for MENDs (red), isotropic magnetoelectric nanoparticles (MENPs, black), and CFONDs (grey). **i,** α*_ME_* for MENDs as a function of AMF frequency at *H*_AMF_ = 10 mT and OMF magnitude 220 mT. **j,** α*_ME_* for MENDs as a function of AMF amplitude for ƒ_AMF_ =150 Hz and OMF magnitude 220 mT.

## Results

### Synthesis and Characterization of Magnetoelectric Nanodiscs

ME modulation relies on effective transduction of an applied MF into a time-varying electric field. Although multiferroic materials can exhibit direct magnetoelectricity (e.g. Cr_2_O_3,_ BiFeO_3_), the effect is enhanced by orders of magnitude via strain-mediated coupling at the interface between a magnetostrictive (MF to strain, e.g. Metglas alloys, Terfenol-D or CoFe_2_O_4_) and a piezoelectric material (strain to electric polarization, e.g. lead zirconate titanate (PZT) or BaTiO_3_)^34^. Core-shell architectures are advantageous in ensemble applications of ME nanomaterials, as the separation between the more conductive magnetostrictive cores prevents leakage of charge generated in each piezoelectric shell^35^. Prior reports on modulation of biological signaling via ME nanomaterials relied on isotropic spherical magnetostrictive cores coupled to piezoelectric shells ^27,28^. Magnetostriction scales with magnetic anisotropy, which is a combination of magnetocrystalline anisotropy and the shape anisotropy of the particles ^36,37^. Consequently, we hypothesized that anisotropic nanodiscs would exhibit greater ME coefficients than the spherical counterparts. We computationally and experimentally tested this hypothesis in a common biochemically inert materials system of Fe_3_O_4_, CoFe_3_O_4_, and BaTiO_3_^38–40^.

Despite the cubic symmetry of the crystal structure, hexagonal magnetite nanodiscs (MNDs) could be synthesized through reduction of hexagonal hematite (Fe_2_O_3_) via previously established protocols (**Fig. 1b**, Fig. S1,2, Methods)^17,41,42^. MNDs with 231±29 nm diameter and 31±9nm thickness were then coated with a CoFe_3_O_4_ shell via an organo-metallic method, resulting in Fe_3_O_4_-CoFe_3_O_4_ core-shell nanodiscs (CFONDs, 240±42 nm diameter **Fig. 1c**, Fig. S2,3, Methods). Piezoelectric BaTiO_3_ shells were then grown onto the CFONDs via a combination of sol-gel synthesis and calcination yielding core-double shell Fe_3_O_4_-CoFe_2_O_4_-BaTiO_3_ ME nanodiscs (MENDs, 250±41 nm diameter, **Fig. 1d**, Fig. S2,4, Methods).

The addition of the CoFe_2_O_4_ shell was motivated by the greater magnetocrystalline anisotropy and magnetostrictive response of this material as compared to Fe_3_O_4_. In addition to a greater bulk magnetostriction coefficient (250 ppm for CoFe_2_O_4_ vs. 25 ppm for Fe_3_O_4_), the interfacial spring exchange effect increases the anisotropy and thus the magnetostrictive response of the core-shell particles^22,43^. Additionally, it has been shown that direct interfacing of materials with different magnetostrictive constants yields an enhanced magnetically-induced strain^44^. To test this, the magnetoelectric coefficient (α*_ME_*) of MENDs was first numerically predicted and then determined experimentally (**Fig. 1e**, Fig. S5).

Using micromagnetic simulation to quantify the effective deformation of nanoparticles due to magnetostriction, we compared CFONDs with spherical Fe_3_O_4_-CoFe_2_O_4_ core-shell nanoparticles of the same volume and volumetric ratio of CoFe_2_O_4_ to Fe_3_O_4_ (Methods). We found that the increase in shape anisotropy in CFONDs affords three orders of magnitude enhancement in effective deformation over spherical nanoparticles (0.83026 ppm vs. 0.000808 ppm) (**Fig. 1f**). For the accuracy of the micromagnetic simulation, the remnant state of CFONDs was found to be either a magnetic vortex or an in-plane macrospin as observed with magnetic force microscopy (Fig. S6). The interaction between Fe_3_O_4_ and CoFe_2_O_4_ in CFONDs yields almost 40 times enhancement in effective deformation as compared to Fe_3_O_4_ MNDs with the same volume (0.01898 ppm, **Fig. 1f**) due to the interfacial spring exchange coupling and surface magnetostriction. The results of the micromagnetic simulation were then introduced into a finite element model of the piezoelectric response of the BaTiO_3_ shell, demonstrating the scaling of the magnetoelectric coefficient (α*_ME_*) with the magnetostriction effect. Together, our numerical simulations predict improved ME coupling in core-double shell MENDs as compared to MNDs or isotropic ME nanoparticles of the same volume (**Fig. 1g**, Fig. S7).

To experimentally evaluate α*_ME_*of the MENDs we expanded on the prior work that leveraged electrochemical means to determine ME coupling from nanoparticles dispersed in an electrolyte^45^ and applied this approach to MEND films^46–49^. Using Tyrode’s solution as an electrolyte, we constructed a three-electrode electrochemical cell, where the working electrode was comprised of a MEND layer on a conductive indium tin oxide substrate with platinum (Pt) and silver/ silver chloride (Ag/AgCl) electrodes serving as counter and reference, respectively (**Fig. 1e**, Fig. S5). To observe ME behavior, the nanoparticles were simultaneously exposed to a constant offset magnetic field (OMF), which magnetized them to near saturation where the change in slope of the magnetization as a function of field amplitude is maximized^50^, and an alternating magnetic field (AMF) which drove magnetization change to induce magnetostriction. The potential change under the different amplitudes and frequencies of MFs was recorded at a constant current of ∼10 nA, allowing us to measure the electric double layer capacitance, while avoiding faradaic reactions. The potential changes were normalized to the AMF amplitude and the MEND film thickness to obtain the α*_ME_* values. The α*_ME_* of MENDs was measured to be 150 mV mT^−1^cm^−1^, which is 4 times higher than the peak value for isotropic CoFe_2_O_4_-BaTiO_3_ core-shell MENPs (**Fig. 1h**, Fig. S8). The discrepancy between the simulated >1000 times increase in the strain response (**Fig. 1f**) and the experimentally observed 4 times increase in α*_ME_* can likely be attributed to the compensation of the electric polarization in a multi-domain piezoelectric shell (Fig. S9) and the lattice mismatch between the cubic CoFe_2_O_4_ and the tetragonal BaTiO_3_ resulting in strain relaxation at the interface, which highlights the opportunity for further materials optimization.

The ME coefficient peaked at 220 mT of OMF, which is near the saturation field of the MENDs recorded via vibrating sample magnetometry (Fig. S10). Fixing the amplitude of the OMF at 220 mT, the α_ME_ was then found to increase with increasing AMF frequency (**Fig. 1i**, Fig. S11a), which is expected for frequencies below the piezoelectric resonance in the MHz range^51–53^. The AC magnetic field amplitude did not have an impact on the α*_ME_* (**Fig. 1j**, Fig. S11b). The ME measurement approach was independent of the identity or concentration of ions in the electrolyte, as α*_ME_* was found to be the same in normal and 10-times concentrated phosphate buffered saline (PBS) as in Tyrode’s solution (Fig. S11c) despite the differences in voltammograms expected from the variation in double-layer capacitance (Fig. S11d).

### MEND-Mediated Neuromodulation In Vitro

We first evaluated the efficacy of ME modulation mediated by MENDs in cultured primary hippocampal neurons virally transduced to express a fluorescent calcium (Ca^2+^) indicator GCaMP6s as a proxy for neuronal activity (Methods). ME neuromodulation with MENDs was expected to be most effective when the particles were in direct contact with neuronal membranes. Consequently, all experiments in vitro were conducted following a 1 h incubation period to allow the particles to precipitate onto the cells. The MEND density on the cell membranes was measured by quantifying their mass on each sample and calculating the area occupied by the neuronal network. Robust increases in GCaMP6s fluorescence were observed in neurons upon exposure to 10 s of combined 220 mT OMF and 10 mT, 1 kHz AMF in the presence of MENDs delivered at a density of 1 µg/mm^2^ (**Fig. 2a, b**, Supporting Video 1). This excitation could be repeated two times, and a decreased response was observed following the third MF epoch potentially due to excitotoxicity, indicating the need for optimization to achieve reliable neuromodulation with MENDs.

**Fig. 2.**
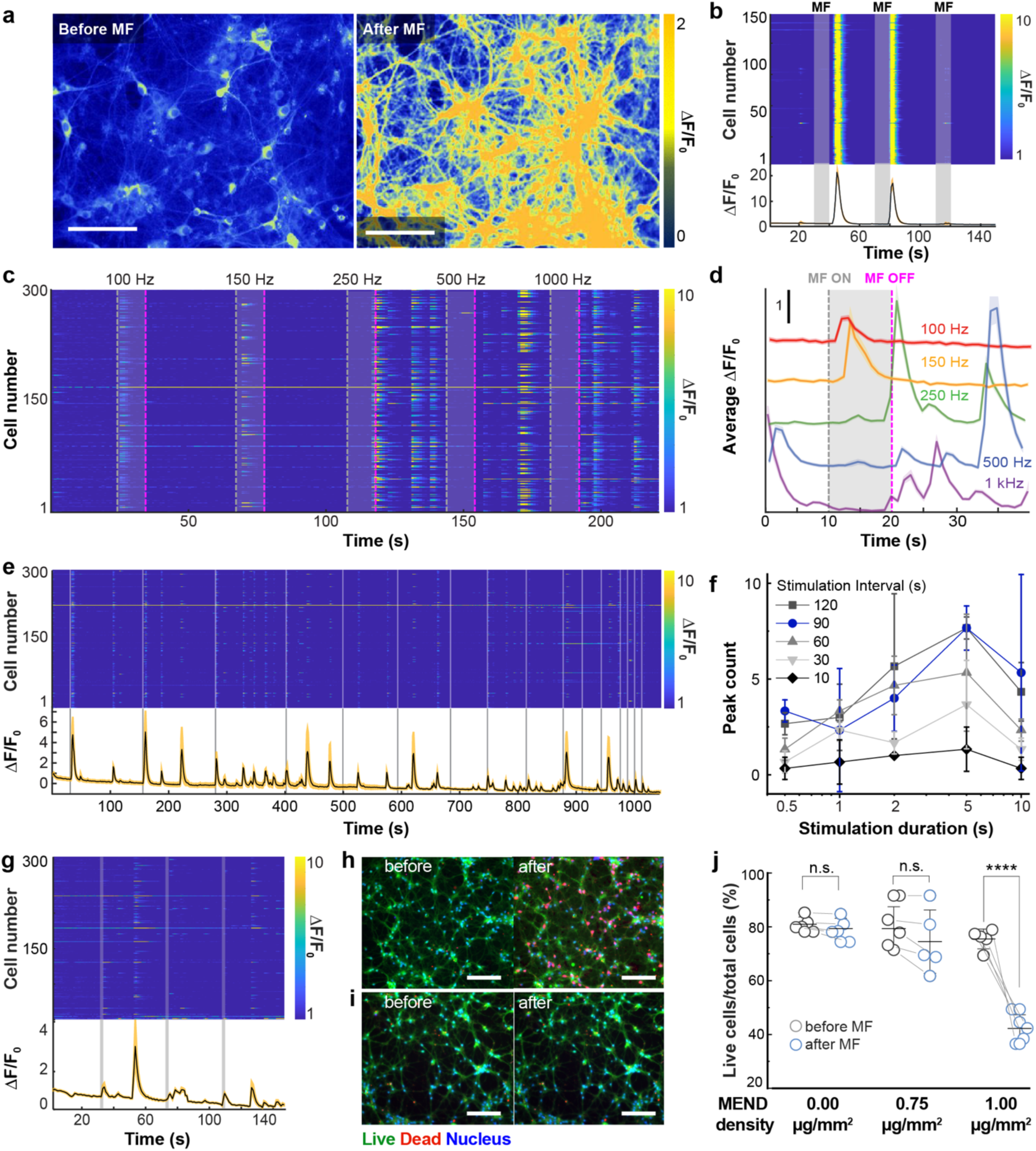
MEND-mediated neuronal stimulation in vitro. a, Relative GCaMP6s fluorescence change (ΔF/F_0_) in hippocampal neurons decorated with MENDs before (left) and after (right) magnetic field application. Scale bars: 150 µm. **b,** Individual (top) and average (bottom) of the relative GCaMP6s fluorescence change (ΔF/F_0_) of the hippocampal neurons in response to 10 mT AMF with 1 kHz frequency and 220 mT OFM. **c,** Individual and **d,** average traces of GCaMP6s ΔF/F_0_ in 300 hippocampal neurons decorated with MENDs in response to 10 mT AMF with frequencies 100, 150, 250, 500, and 1000 Hz (OMF magnitude 220 mT). The grey and magenta dashed lines indicate the beginning and end of MF stimulation, respectively. **e,** Individual cell (top) and mean (bottom) GCaMP6s fluorescence changes in 300 neurons in response to 2 s MF epochs applied at varying intervals (OMF 220 mT; AMF 150 Hz, 10 mT). **f,** Number of GCaMP6s fluorescence peaks as a function of stimulation epoch length for rest intervals of 10, 30, 60, 90, and 120 s (OMF 220 mT; AMF 150 Hz, 10 mT). **g,** Individual cell (top) and mean (bottom) GCaMP6s fluorescence changes in response to 2 s MF epochs at 40 s intervals for 0.75 µg mm^−2^ MEND density. **h, i,** Fluorescent image of a Live-Dead assay in neurons before and after 3 cycles of MF at **h,** 1 µg mm^−2^ and **i,** 0.75 µg mm^−2^ MEND densities. **j,** The change in live cell ratio (normalized to the total number of the cells counted with Hoest staining) following 3 cycles of MF for neurons decorated with different MEND densities. Statistical significance was tested via one-way ANOVA and Tukey’s multiple comparison tests (n = 5 plates per condition, P=3.79×10^−7^ for 1 µg mm^−2^; P=0.79 for 0.794 µg mm^−2^; P=0.998 for 0.75 µg mm^−2^; ****P≤0.0001, n.s. P>0.05).

Although α*_ME_*scales with the AMF frequency, resulting in larger fluctuations of the membrane potential, high frequency (>150 Hz) stimulation yields neuronal conduction block^54,55^. This suggests the existence of an optimal frequency range for neuromodulation mediated by MENDs. Indeed, for AMF frequencies >150 Hz, the neurons appeared silenced during MF (220 mT OMF and 10 mT AMF) exposure, and rebound activity was observed upon MF removal (**Fig. 2 c,d**, Fig. S12, Supporting Videos 2 and 3). In contrast, at frequencies of 100 Hz and 150 Hz, GCaMP6s fluorescence transients emerged during the MF (Fig. S12). Furthermore, at frequencies <100 Hz, probability of observing GCaMP6s transients in response to MF stimuli was reduced (Fig. S12), which could in part be due to the reduced α*_ME_* (Fig. 1i). Therefore, 150 Hz was chosen as the AMF frequency for the following experiments in vitro and in vivo.

We further explored neuronal excitation as a function of stimulation duration and interval between field epochs. We quantified the number and latency of GCaMP6s fluorescence transients for field epochs (220 mT OMF; 10 mT, 150 Hz AMF) ranging between 0.5-10 s separated by rest epochs between 10-120 s. Stimulation epochs longer than 2 s produced a greater number of GCaMP6s transients than 1 s and 0.5 s stimulation epochs regardless of the rest epoch length (**Fig. 2e, f** and Figure S13, Supporting Video 4). However, unlike short stimulation epochs that produced 2.0 ± 1.6 GCaMP6s fluorescence transients per epoch, longer stimulation epochs yielded unsynchronized responses (3.9±3.09 GCaMP6s transients, **Fig. 2f**). We also observed a decrease in the number of magnetically evoked GCaMP6s transients for stimulation epochs separated by ≤90 s. This is consistent with activity suppression observed in mature hippocampal cultures between bursts occurring spontaneously every 25-83 s.^56^

As expected, the extent of neuronal response to the MF stimuli (220 mT OMF; 10 mT, 150 Hz AMF) scaled with the MEND density. At MENDs densities of 1 µg/mm^2^, 0.75 µg/mm^2^, 0.5 µg/mm^2^, and 0.25 µg/mm^2^, 73.9±15.2%, 74.1±23.6%, 46.3±27.6%, and 27.6±23.3% of neurons were responsive to MF, respectively (**Fig. 2g**, Fig. S14a,b). In the absence of MENDs, no significant response to identical magnetic field conditions was observed (Fig. S14c). The excitotoxity of the ME modulation was similarly evaluated at different MEND densities. The viability of neurons prior to ME modulation was not affected by the presence of MENDs alone at 1 µg/mm^2^ or 0.75 µg/mm^2^ densities as compared to cultures without MENDs (Fig. S15). At a MEND density of 1 µg/mm^2^, three 10 s repetitions of combined MF stimulus led to a decrease in neuronal viability in vitro from 73.4±3.65% to 40.1±5.09%, while no significant decrease in viability was observed at a MEND density of 0.75 µg/mm^2^ (**Fig. 2h-j**, Fig. S15). The observed excitotoxicity of ME neuromodulation at a MEND density of 1 µg/mm^2^ explained the reduced cell responsiveness to the third stimulation epoch at this MEND density (Fig. 2a, b, Supporting Video 1). Additionally, these findings motivated the use of lower MEND concentrations and AMF frequencies to achieve repeatable and safe ME neuromodulation.

Combined MF conditions employed for ME neuromodulation (220 mT OMF; 10 mT, 150 Hz AMF) were not anticipated to evoke mechanical or thermal effects on neurons due to the inability of the MENDs to oscillate in the presence of a near-saturation OMF or dissipate observable heat at AMF frequencies below the kilohertz range. Indeed, the combined MF conditions did not evoke excitation in neurons decorated with CFONDs that constitute magnetic cores of MENDs even in the presence of virally delivered thermosensitive (transient receptor potential vanilloid receptor 1, TRPV1, Fig. S16a, b) or mechanosensitive (TRPV4, Fig. S16c, d) ion channels.

### Mechanistic Investigation of MEND-Mediated Neuromodulation

The potential generated by individual MENDs estimated from α*_ME_* measurements at the field conditions employed in our in vitro studies is ∼35 μV, which is significantly below the neuronal activation threshold (15-30 mV)^33^. Consequently, we sought to investigate the mechanisms underlying the observed neuronal responses. We hypothesized that MENDs exert repetitive subthreshold stimulation on neuronal membranes, which is then integrated across multiple periods of the AMF resulting in neuronal depolarization facilitation (**Fig. 3a**, Supplementary Note S1).

**Fig. 3.**
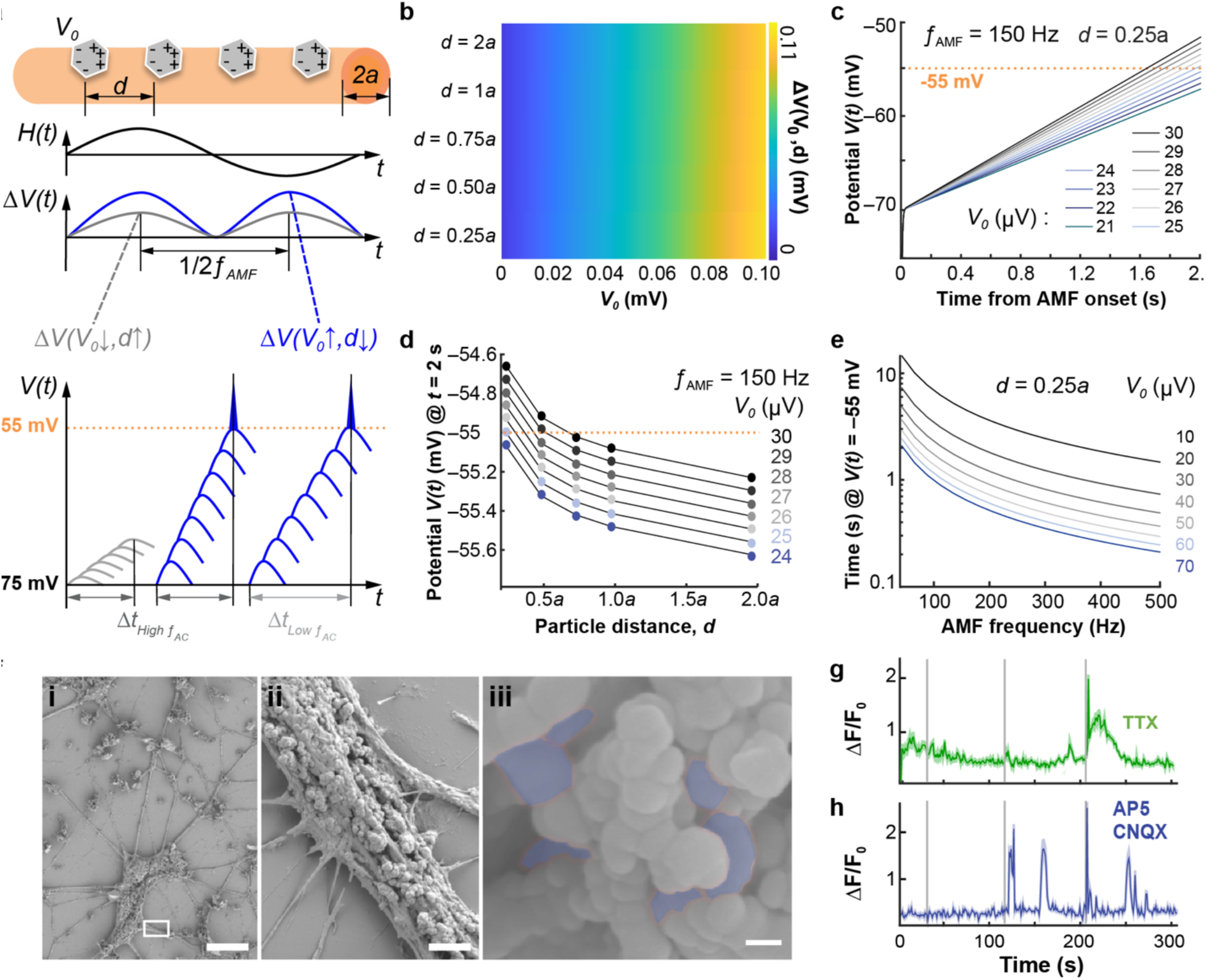
| Mechanistic study of MEND-mediated neuromodulation. a, Illustration of stimulation mechanism combining cable model and repeated excitation of neural activity, where *d* is spacing between MEND particles, *a* is cell radius, Δ*V* is the change in membrane potential per half-period of an AMF, and *V*_0_ is the voltage generated by a single MEND. **b,** Calculated membrane potential change Δ*V* for every half-period of an AMF, as a function of *V*_0_ and *d*. **c,** Simulated membrane potential *V*(*t*) after AMF onset to activate MENDs with *V*_0_ = 0.03 mV for varying *a* values. The threshold potential for action potential firing, –55 mV, is indicated with the dashed line. **d,** Simulated membrane potential at a time *t* = 2 s after onset of magnetic field as a function of *d*, for varying *V*_0_ values. *V*_0_ =0.03 mV is the measured value for MEND particles in this study, highlighted in red. **e,** Time to reach threshold membrane potential (–55 mV) from the resting potential (–75 mV) as a function of AMF frequency ƒ_AMF_ for varying *V*_0_ values. **f,** (**i-iii**) SEM images showing MENDs decorating cultured hippocampal neurons. **(ii)** A higher magnification image of the area marked by a box in panel (i). **(iii)** MENDs on the neuron surface shaded in blue as identified by the presence of iron, titanium, and barium in energy dispersive X-ray spectroscopy. Scale bars: 20 µm (i), 5 µm (ii), 100 nm (iii). **g, h,** GCaMP6s fluorescence changes in neurons decorated with MEND following 2 s stimulations (OMF 220 mT; AMF 150 Hz, 10 mT, marked by vertical grey bars) in the presence of **g,** tetrodotoxin (TTX, 1 µM, green) or **h,** a cocktail of (2R)-amino-5-phosphonovaleric acid (AP5, 100 µM) and 6-cyano-7-nitroquinoxaline-2,3-dione (CNQX, 20 µM) (blue).

To simulate neuronal facilitation during ME stimulation mediated by MENDs, a model of repetitive neuronal excitation^57^ was combined with the steady-state solution for the three-dimensional (3D) cable theory^58^ (Supplementary Note S1, Figure S17). For a single period of AMF, the integrated change in membrane voltage Δ*V* was calculated using cable theory as a function of a single MEND potential *V*_0_ and the average intra-particle distance, *d* (**Fig. 3b**). Δ*V* depended more strongly on V_0_ than on the inter-particle distance. Only 10% increase in Δ*V* could be achieved by an 8-times decrease in the particle spacing. The change in membrane potential was then superimposed onto the prior state of the membrane potential, which depended on the stimulation history, stochastic fluctuations, and the duration of the refractory period and hyperpolarization delay for a given neuron (Supplementary Note S1). **Fig. 3c** describes the change in membrane potential over time produced by the AMF oscillating at 150 Hz for MENDs with *d* = 0.25*a* (*a* is axonal radius) and various values of *V*_0_. These findings indicate that neuronal potential (initially set to –70 mV) can reach threshold values of –55 mV in 2 s if the MENDs are spaced by 0.25*a* and generate *V*_0_ >24 µV on the membrane, which is consistent with our experimental observations (Fig. 2d-f).

Our model employs the steady-state solution for the cable equation and temporal summation of the membrane potential fluctuations. This approximation was motivated by the same order of magnitude of the duration of AMF pulses (6.67 ms at 150 Hz) and the neuronal time constant (1-20 msec)^59,60^ At lower frequencies (<100 Hz), AMF pulse length precludes the superposition of generated potentials Δ*V*, which may further explain lower rates of neuronal responses at these frequencies (Fig. S12). As our model does not recapitulate the neuronal conduction block, the latency to reach the threshold potential monotonically decreases with increasing *V*_0_ (**Fig. 3d**) and AMF frequency (**Fig. 3e**), which contrasts with our experimental observations at AMFs with frequencies >150 Hz. Another assumption of our model was the uniform distribution of MENDs on neuronal membranes. To test the validity of this assumption, scanning electron microscopy (SEM) was applied to the MEND-decorated neurons (**Fig. 3f**). Since synaptic boutons in intact neurons (Fig. S18) exhibit dimensions comparable to MENDs, energy-dispersive X-ray spectroscopy mapping (Fig. S19) was employed to distinguish the particles from neuronal structures by their element content, revealing the presence of individual MENDs as well as few-particle aggregates on the cell membranes.

Consistent with our model, neuronal depolarization, albeit diminished, was observed even in the presence of the voltage-gated sodium channel blocker tetrodotoxin (TTX, 1 µM), as the threshold potential could be reached regardless of the state of the sodium channels (**Figure 3g**, Figure S20a, Supporting Video 5) at the density of 0.75 µg/mm^2^ MENDs. Moreover, MEND-mediated excitation did not rely on synaptic transmission within the neuronal network, as GCaMP6s transients were observed in response to a MF (220 mT OMF; 10 mT, 150 Hz AMF) in the presence of a cocktail of the α-amino-3-hydroxy-5-methyl-4-isoxazolepropionic acid (AMPA) receptor antagonist ((2R)-amino-5-phosphonovaleric acid, AP5, 100 µM) and the non-N-methyl-D-aspartate (non-NMDA) receptor antagonist (6-cyano-7-nitroquinoxaline-2,3-dione, CNQX, 20 µM) (**Figure 3h**, Figure S20b, Supporting Video 6).

Although our model offers qualitative description of possible mechanisms underlying MEND-mediated neural excitation, it likely underestimates the magnitude of the effect as it does not consider the spatial summation of the membrane potential fluctuations ^61^ or the diffuse-charge accumulation of ions around MENDs^62,63^. A detailed model that includes the kinetics and distribution of specific ion channels, neuronal geometry, external excitatory and inhibitory inputs, as well as the effects of the MEND polarization on the surrounding ion distribution will most likely yield a closer quantitative agreement with experiments^64,65^. Nevertheless, the framework proposed here informs conditions for MEND-mediated modulation in vivo.

### MEND-Mediated Neuromodulation In Vivo

We next evaluated whether MENDs could mediate ME neuromodulation in deep-brain structures in mice (**Fig 4a**). MENDs (1.5 µL volume, 1.5 mg/mL or a control solution) were injected into the left ventral tegmental area (VTA), a predominantly dopaminergic structure critical to reward processing ^66,67^, in wild-type BL6/57 mice (n=6 for every sample group). Following a 1-week recovery period, the mice were subjected to 3 cycles of MF stimulation (220 mT OMF, 10 mT, 150 Hz AMF; 2 s epochs separated by 90 s rest periods), and their brain tissue was analyzed with respect to expression of an immediate early gene c-Fos, a marker of neuronal activity (Methods).^68^ Consistent with our analyses in vitro, the percentage of c-Fos expressing cells relative to all cells in a field of view marked by nuclear stain 4’,6-diamidino-2-phenylindole (DAPI) was significantly higher in the VTA of mice injected with MENDs and exposed to the MF in comparison to controls injected with phosphate buffered saline (PBS), MNDs without the piezoelectric BaTiO_3_ shell (1.5 µL at 1.5 mg/mL), or mice injected with MENDs and not exposed to the MF (**Fig. 4b,c**, Fig. S21). The expression of c-Fos in response to MF stimulation was also significantly higher in the nucleus accumbens (NAc) and the medial prefrontal cortex (mPFC), the excitatory targets of the VTA^66,67^, in mice injected with MENDs than in controls (**Fig. 4d-g**, Figures S22, S23). Notably, reducing the MEND concentration to values comparable to the experiments in vitro (1.5 µL at 0.5 mg/mL), resulted in c-Fos expression levels in the VTA, NAc, and mPFC similar to those found with higher particle concentrations (**Fig. 4b-g**, Figure S21-S23).

**Fig. 4.**
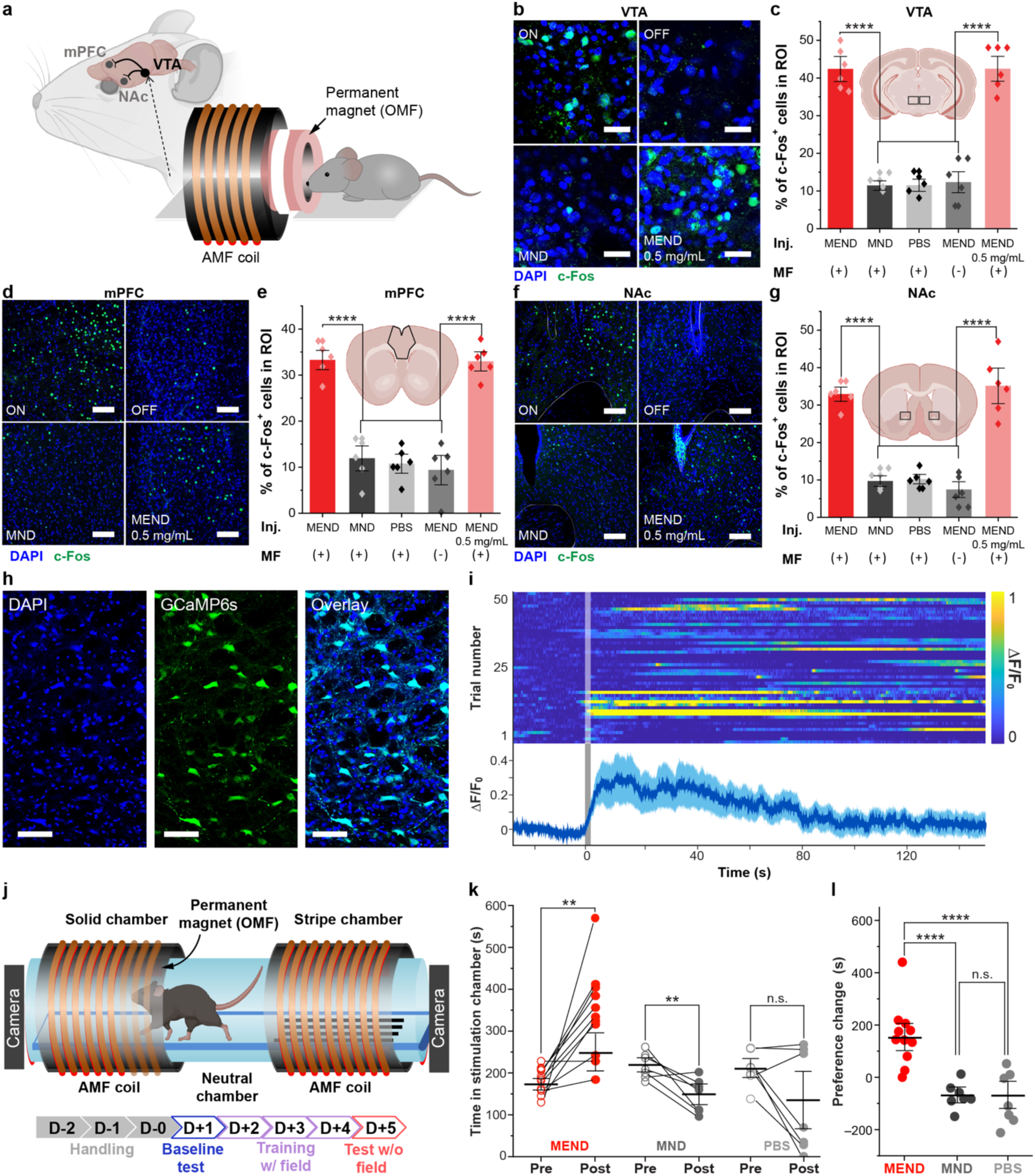
| MEND-mediated neuronal stimulation in mice. a, Schematic of MEND-mediated neuromodulation. MEND particles were injected into the mouse ventral tegmental area (VTA), and the mice were placed inside a permanent magnet field providing OMF of 220 mT and a surrounding solenoid providing AMF with an amplitude 10 mT and a frequency 150 Hz. **b,** Confocal images of c-Fos-expressing neurons among DAPI-marked cells in the VTA. **Top left:** MENDs (1.5 mg/ml) with (+) magnetic stimulation (ON); **top right:** MEND particles (1.5 mg/ml) without (–) magnetic stimulation (OFF); **bottom left:** control MNDs (1.5 mg/ml) + magnetic stimulation; **bottom right:** MENDs (0.5 mg/ml) + magnetic stimulation. Scale bars are 25 µm. **c,** Quantification of c-Fos positive neurons for the conditions shown in (B) as well as the subjects injected with PBS and exposed to magnetic field. **d,** Confocal images and **e,** quantification of c-Fos positive neurons in the medial prefrontal cortex (mPFC) for the same conditions shown in (B) and (C). **f,** Confocal images and **g,** quantification of c-Fos positive neurons in the nucleus accumbens (NAc) for the same conditions shown in **b** and **c**. **d, f,** Scale bars are 100 µm. **c, e, g,** Statistical significance was tested via one-way ANOVA and Tukey’s multiple comparison tests (n = 6 mice per condition, VTA F_3,20_=93.2 P=1.21×10^−12^; mPFC F_3,20_=60.81 P=1.21×10^−11^; VTA F_3,20_=62.03 P=1.29×10^−12^; ****P≤0.0001). **h,** Representative confocal images showing expression of GCaMP6s in the VTA. Scale bar: 50 µm. **i** Fiber photometry recordings of relative GCaMP6s fluorescence change (ΔF/F_0_) in the VTA of anaesthetized mice injected with MENDs in the same region. (Top) Individual trial ΔF/F_0_. (Bottom) Average ΔF/F_0_ across trials shown above. Solid line represents mean and shaded areas mark s.e.m. (n=4 mice, 10-15 trials per mouse). The grey square box indicates magnetic field epochs (2s, OMF 220 mT, AMF 10 mT, 150 Hz). **j** Schematic of the place preference behavioral apparatus (top) and experimental timeline (bottom). **k,** Time spent in the stimulation chamber out of a total test time of 600 s, for pre-(Day 1, open markers) and post-(Day 5, solid markers) learning. Paired t-test was performed for MEND (n=11) and MND (n=7) groups, and Wilcoxon signed-rank test was performed for PBS (n=7) group because the data did not follow normal distribution. ***P ≤ 0.001, **P ≤ 0.01, *P ≤ 0.05, n.s. P>0.05). **l,** The change in time spent in stimulation chamber between Day 1 and Day 5. (One-way ANOVA with Tukey’s post-hoc comparison test, ***P ≤ 0.001, **P ≤ 0.01, *P ≤ 0.05, n.s. P>0.05.)

Next, we employed fiber photometry in the VTA to record neural activity during ME modulation in mice (Fig. S24). We injected MEND (1.3 µL, 1.5 mg/mL) solution (or control MNDs or PBS) into the VTA of wild type mice, along with the adeno-associated virus carrying GCaMP6s under a pan-neuronal human synapsin promoter (AAV9-hSyn::GCaMP6s, 0.3 µL), which was followed by an implantation of a 200 µm silica optical fiber into the same location. Fluorescent imaging confirmed GCaMP6s expression in cell bodies in the VTA, and in axons projecting to NAc and mPFC (**Fig. 4h**, Fig. S25). Following a 2-week incubation period, GCaMP6s fluorescence increase was observed in response to 2 s MF stimulation (220 mT OMF, 10 mT, 150 Hz AMF) in mice injected with MENDs but not in those injected with control particles or PBS (**Fig. 4i**, Fig. S26).

Based on prior work on dopamine-dependent reward circuits^67,69^, we tested whether ME stimulation in the VTA could induce place preference in a three-chamber assay in mice. To enable neural excitation mediated by MENDs, we designed a custom place preference arena comprised of two stimulation chambers with different floor patterns (stripes vs. solid) equipped with identical electromagnets to supply the AMF and permanent magnets supplying the OMF separated by a neutral chamber (**Fig. 4j** and Fig. S27). Following a 1-week recovery period, mice injected with MENDs (1.5 µL, 1.5 mg/mL), control MNDs, or PBS into the VTA were habituated to the arena for 3 days, and their baseline preference was then tested for 10 min on day 1 (**Fig. 4j**). During learning days 2-4, the mice were allowed to explore the entire arena for 10 min while MF (220 mT OMF, 10 mT, 150 Hz AMF, 2 s epochs separated by 90 s rest periods) was applied in their baseline non-preferred chamber. On test day 5, the mice explored the entire arena in the absence of MF stimuli in either chamber. On days 2-4 learning period, mice injected with MENDs in the VTA exhibited an increased tendency toward the magnetic stimulation chamber, as compared to the mice injected with control solutions (Fig. S28). On day 5 (Test), mice injected with MENDs demonstrated a conditioned preference to the stimulation chamber (as compared to Day 1, Baseline), while mice injected with MND and PBS, respectively, exhibited or trended toward aversion for the stimulation chamber (**Fig. 4k**). The average preference change for the stimulation chamber in MEND-injected mice was significantly greater and positive, while it was slightly negative in mice injected with control solutions (**Fig. 4l**). This aversion can likely be attributed to the minor vibrations and noise associated with electromagnets operating at 150 Hz.^70,71^

We next assessed the biocompatibility of MEND-mediated neuromodulation. We hypothesized that wireless neuromodulation with MENDs would have reduced tissue damage when compared to electrode implantation.^72,73^ To test this hypothesis, we quantified microglial activation marker (ionized calcium-binding adaptor molecule1, Iba1), astrocytic glial fibrillary acidic protein (GFAP), and macrophage activation marker (cluster of differentiation 68, CD68) surrounding MENDs relative to PBS injection controls and to 300 µm stainless steel microwire implants commonly used for electrical neuromodulation in mice. We observed negligible increases in Iba1, GFAP, and CD68 expression in mice injected with MENDs compared to those injected with PBS. By contrast, all markers were significantly upregulated near the microwire implants relative to MEND injections (Fig. S29, S30).

## Conclusion

We colloidally synthesized anisotropic magnetoelectric nanodiscs (MENDs) with core-double shell architecture (Fe_3_O_4_-CoFe_2_O_4_-BaTiO_3_), where layers of magnetostrictive CoFe_2_O_4_ and piezoelectric BaTiO_3_ were grown on Fe_3_O_4_ templates. The MENDs exhibited enhanced ME coefficient as compared to previously reported isotropic CoFe_2_O_4_-BaTiO_3_ nanoparticles. When decorated onto neuronal membranes in vitro at densities 0.5-1 µg/mm^2^, MENDs mediated robust and rapid excitation with combined OMF (220 mT) and AMF (10 mT) at frequencies up to 150 Hz. Given the theoretical limits on the electrical potential that can be generated by nanometer-scale particles, we investigated the biophysical mechanisms that could underly ME neuromodulation. By combining cable theory with a model of repeated neuronal depolarization, we have shown that voltages generated by individual MENDs can be integrated across the periods of the driving AMF and across multiple particles. We found that the change in membrane potential increases with increasing AMF frequency commensurate with the scaling of the ME coefficient, however our model did not recapitulate the high-frequency conduction block. Additionally, decreasing the inter-particle spacing on neuronal membranes reduced the latency for potential threshold crossing and action potential firing. Guided by these insights, we translated MEND-mediated stimulation to experiments in vivo, achieving remote magnetic control of neuronal activity in the VTA of wild-type mice at particle concentrations as low as 0.5 mg/mL (1.5 µL injections). Robust neural excitation in the VTA mediated by MENDs enabled remote magnetic control of behavior, driving conditioned place preference in genetically intact mice, making this approach a minimally invasive alternative to electrical DBS. We anticipate that the materials and biophysical insights developed in this work will set the stage for broad application of minimally invasive ME modulation in neuroscience. MEND technology may uniquely empower applications where implementation of genetic neuromodulation methods or invasive hardware is challenging, such as experiments in translational non-rodent models or in peripheral nervous system targets.

## Acknowledgements

This work was funded in part by the Pioneer Award from the National Institutes of Health and National Institute for Complementary and Integrative Health (DP1-AT011991), NIH BRAIN Initiative and the National Institute for Neurological Disorders and Stroke (R01-NS115576), McGovern Institute for Brain Research at MIT, and K. Lisa Yang and Hock E. Tan Center for Molecular Therapeutics at MIT. Y.J.K. is a recipient of the Mathworks Fellowship. A.T. is a recipient of the National Science Foundation Graduate Research Fellowship and the Paul and Daisy Soros Fellowship for New Americans. The authors are grateful to Prof. James LeBeau and Prof. Caroline Ross in the Department of Materials Science and Engineering for the expertise and resources in electron microscopy and insights into multiferroic properties of composite nanomaterials, respectively; to Prof. Mark Harnett, Prof. Ila Fiete, Enrique Toloza, and Mikail Khona at the McGovern Institute for Brain Research for their ideas and thoughtful feedback on modeling of neuronal activation with magnetoelectric nanodiscs; to Dr. Michael Christiansen at the Swiss Federal Institute of Technology (ETH Zurich) for his feedback on all aspects of our manuscript.

## Author contributions

Y.J.K. and P.A. designed the study. Y.J.K. synthesized and characterized the materials, created a custom electrochemical cell for magnetoelectric effect measurements, performed all in vitro and in vivo experiments, and developed the model for neuronal depolarization. Y.J.K. and F.K. constructed the magnetic field apparatuses for in situ, in vitro, and in vivo experiments, with design input from N.K. and A.B.S. N.K. and Y.J.K. performed micromagnetic and finite element modeling, and magnetic force microscopy. N.K. performed the vibrating sample magnetometry. Y.J.K. and N.D. performed photometry studies and analyses. Y.J.K., E.V. and M.M. performed behavioral assays. Y.J.K., A.T. and E.V. performed biocompatibility studies. Y.J.K., N.D., N.K., A.B.S. and P.A. have analyzed the data. Y.J.K. and N.D. have created figures for the manuscript. All authors have contributed to the writing of the manuscript.

## Data and materials availability

All data needed to evaluate the conclusions in the paper are present in the paper or the supplementary materials. Reasonable quantities of physical samples of the described magnetoelectric nanodiscs are available upon request.

## Supplementary Materials

Methods

Supplementary Note S1

Figures S1-S26

Tables S1-S5

Supporting Videos 1-6

References 1-9

## Materials and Methods

### Synthesis of magnetoelectric nanodiscs

Synthetic procedures for magnetoelectric nanodiscs were reproduced across two institutions (Massachusetts Institute of Technology and Friedrich-Alexander University of Erlangen – Nuremberg). The Fe_3_O_4_ magnetic nanodiscs (MNDs) were synthesized by reducing hematite nanodiscs. Hematite nanodiscs were first produced by heating a uniform mixture of 0.273 g of FeCl_3_·6H_2_O (Fluka), 10 mL Ethanol, and 600 μL of deionized (DI) water in a sealed Teflon-lined steel vessel at 180℃ for 18 hours. After washing the red hematite nanodiscs with DI water and ethanol 3-5 times, the dried hematite was dispersed in 20 mL of trioctyl-amine (Sigma-Aldrich) and 1g of oleic acid (Alfa Aesar/ Thermo Fisher Scientific). For the reduction of hematite to magnetite, the mixture was transferred into a three-neck flask connected to a Schlenk line, and evacuated for 20 min at room temperature, and then heated to 370 ℃ (20 ℃/min) in H_2_ (5%) and N_2_ (95%) atmosphere for 30 min.

The core-shell Fe_3_O_4_-CoFe_2_O_4_ nanodiscs (CFOND) were formed by nucleation and growth of a CoFe_2_O_4_ layer on the surface of MNDs. For this procedure, 120 mg of MNDs (cores) were dispersed uniformly in a precursor solution of 20 mL diphenyl ether (Aldrich), 1.90 mL oleic acid (Sigma Aldrich), 1.97 mL oleylamine (Aldrich), 257 mg cobalt acetylacetonate (Co(acac)_2_, Aldrich), and 706 mg iron acetyl acetonate (Fe(acac)_3_, Aldrich). A three-neck flask including the solution of MND cores and the shell precursors was connected to a Schlenk line. The solution was evacuated and then heated to 100 °C (7 °C/min) for 30 min in N_2_ atmosphere while magnetically stirring at 400 rpm. After closing the N_2_ line, the temperature was increased to 200 °C (7 °C/min) and maintained for 30 min, and then increased to 230 °C (7 °C/min) and maintained for 30 min. The solution was cooled to room temperature (∼30 min), and the resulting CFONDs were washed with ethanol and n-hexane and subjected to centrifugation at 8000 rpm for 8 min; the washing process was repeated 2-3 times. The thickness of CoFe_2_O_4_ layer is controlled by repeating the organometallic synthesis and washing steps described above. To obtain a 5 nm CoFe_2_O_4_ layer, the synthesis was repeated three times.

The Fe_3_O_4_-CoFe_2_O_4_-BaTiO_3_ magnetoelectric nanodiscs (MENDs) were made by formation of BaTiO_3_ shell on the surface of CFOND via the sol-gel method. A mixture comprising 16 mg of CFONDs dispersed in n-hexane, 30 mL of DI water, 6 mL of ethanol, and 2g of poly(vinylpyrrolidone) (Sigma Aldrich) was sonicated for 20 min, which led to segregation of the oil phase. The oil phase and other insoluble solids were removed with a spatula. The hydrophilic CFOND dispersions were then transferred to a three-neck flask connected to a Schlenk line, and then dried in vacuum at 80 ℃ until amber-colored gel was formed on the bottom of the flask. The gel was re-dispersed in the BaTiO_3_ shell precursor solution which was prepared by mixing 0.5 g citric acid (Sigma Aldrich) and 24 µL titanium isopropoxide (Aldrich) dissolved in 15 mL of ethanol and 0.1 g citric acid and 0.0158 g barium carbonate (Aldrich) dissolved in DI water. The solution of CFONDs and BaTiO_3_ precursors were moved to the three-neck flask connected to the vacuum line and kept at 80 ℃ for 12-14 hours. The powders were then moved to a clean ceramic container and heated at 600 ℃ for 2 hours, 700 ℃ for 2 hours, then 800 ℃ for 1 hour, sequentially. To prevent breaking the BaTiO_3_ shell, the furnace door was kept closed until the temperature slowly cooled down to room temperature. The MENDs were dispersed in Tyrode and PBS before being used for in-vitro and in-vivo experiments.

### Structural and Magnetic Characterization of Magnetic Nanomaterials

Structural imaging of MNDs, CFONDs, and MENDs and energy-dispersive X-ray spectroscopy mapping on MEND-including neurons was performed via Scanning Electron Microscopy (SEM, Zeiss Merlin). Transmission electron microscopy (TEM) imaging and single-particle electron diffraction analysis was performed using a FEI Tecnai G2 Spirit TWIN TEM. The diameter and thickness of MNDs, CFONDs, and MENDs were estimated from the ensemble averages of particles in TEM and SEM images. Powder X-ray diffraction patterns of as-synthesized MNDs, CFONDs, and MENDs were collected by a three-circle diffractometer coupled to a Bruker-AXS Smart Apex charged-coupled device (CCD) detector with graphite-monochromated Mo *K*_$_ radiation (λ = 0.71073 Å), and the data were processed with PANalytical HighScore Plus software. Room-temperature hysteresis curves were generated using the combined superconducting quantum interference device (SQUID) and vibrating sample magnetometer (VSM) mode of a Quantum Design MPMS-3 at 300 K. An Agilent 5100 Inductively Coupled Plasma-Optical Emission Spectrometer (ICP-OES) was used to quantify the elemental concentration for the calculation of saturation magnetization. For ICP-OES analysis, nanoparticles were digested in 37% v/v HCL (Sigma-Aldrich) overnight and diluted in 2 wt % HNO_3_ (Sigma-Aldrich).

### Micromagnetic simulations

Full magnetoelastic simulations were performed with MuMax3 magnetoelastic extension using literature values of magnetic and elastic constants for the materials (**Table S1**) and a cell size of 2×2×2 nm^3^. The simulation was performed at discrete magnetic fields corresponding to the maxima of an oscillating field with an amplitude of 10 mT around an offset at 220 mT. Increasing the strength of the elastic damping constant, a steady-state elastic displacement could be achieved within a simulation time of 72-168 hours running on a GeForce RTX 3060 GPU. The steady state emerged after ∼35 ns following the initial field application, and 10 ns was required for subsequent field steps. Hence, all simulations were performed with the initial field value for 40 ns, and each subsequent field value for 15 ns. To quantify the normalized displacement of each particle at the maxima of an oscillating field with an amplitude of 10 mT around an offset at 220 mT, we averaged root-mean-square displacement normalized to the solid diagonal of each simulation cell. To calculate the deformation of each particle under the oscillating field with an amplitude of 10 mT around an offset at 220 mT, we averaged the difference in normalized displacements at two sequent maxima. Note that due to limits of MuMax3 Magnetoelastic extension we had to use the same sign in the anisotropy of the Fe_3_O_4_ and CoFe_2_O_4_ for the core shell particles. This is an acceptable approximation as a lower bound of the strain in the composite particle. With a difference in anisotropy sign, the very large magnetostriction could not be simulated with our existing computational infrastructure in as the resultant pressure waves took microseconds to dissipate.

**Table S1.** Materials parameters employed in micromagnetic simulations of magnetostriction.

| Constant | $\text{Fe}_3\text{O}_4$ | $\text{CoFe}_2\text{O}_4$ |
| --- | --- | --- |
| Saturation Magnetization | $480 \times 10^3 \text{ A/m}^{74,75}$ | $445 \times 10^3 \text{ A/m}^{76}$ |
| Exchange Coupling Strength | $1.3 \times 10^{-12} \text{ J/m}^{77}$ | $1.7 \times 10^{-11} \text{ J/m}^{76}$ |
| Interlayer exchange coupling | $1 \times 10^{-12} \text{ J/m to Co}^{78}$ | $1 \times 10^{-12} \text{ J/m to Fe}^{78}$ |
| Crystalline Anisotropy Axis | $(1, 1, 1)^{77}$ | $(1, 1, 1)^{79}$ |
| Crystalline Anisotropy constant | $13500 \text{ J/m}^3^{36,77}$ | $240000 \text{ J/m}^3^{36,76}$ |
| Density | $5110 \text{ kg/m}^3$ | $5230 \text{ kg/m}^3$ |
| Magnetoelastic coupling constant | $1.6 \times 10^7 \text{ J/m}^3^{77}$ | $3.2 \times 10^7 \text{ J/m}^3^{80}$ |
| C11, C12, C44 | $240 \times 10^9, 135 \times 10^9, 87 \times 10^9 \text{ N/m}^2^{77}$ | $244 \times 10^9, 142 \times 10^9, 75 \times 10^9 \text{ N/m}^2^{80}$ |

### Design and fabrication of electromagnets

For magnetoelectric coupling coefficient measurements and Ca imaging in vitro, TEMCo 14 AWG copper magnet wire was wound around a C-shaped magnetic core with an 8 mm gap. The coil was connected to a power supply (Crown DC-300A Series II) and signal generator (Picoscope 2204A) to generate a magnetic field combining a static magnetic field offset (OMF, magnitude: 0 – 320 mT) and an alternating magnetic field (AMF, frequency: 0 – 1000 Hz, amplitude: 0 – 14 mT). For in-vivo experiments we separated the apparatuses that generated the OMF and AMF to generate a uniform, time varying, magnetic field over a larger volume. To generate the AMF (10 mT, 150 Hz) for c-Fos expression and fiber photometry experiments, we used a TEMCo 14 AWG copper magnet wire solenoid with a 10 cm inner diameter connected to the power supply and the signal generator. To generate the DC magnetic field (220 mT), we used a ring-shaped permanent magnet (3.81 cm od x 1.905 cm id x 0.3175 cm thick, K&J Magnetics, Inc. RX8C2) that was positioned in the center of the AMF coil. To apply magnetic field during behavior experiments, we designed a custom arena comprised of two-connected cylindrical chambers (FixtureDisplays® clear acryalic tube, 75 mm diameter, 2 mm wall thickness, and 205 mm long) that was divided into two stimulation chambers (11 cm) separated by a neutral area (19 cm) that contained the animal entry port. The entry port was covered with 3D printed acrylic curved plate after the entrance of mouse. To generate OMF during behavioral experiments, two rectangular permanent magnets (7.62 cm x 7.62 cm x 1.27 cm thick, K&J Magnetics, Inc. BZ0Z08-N52) were placed at the top and bottom of each stimulation chamber. To generate AMFs during behavioral experiments, solenoids (TEMCo 14 AWG copper magnet wire) were wound around each stimulation chamber and connected to the power supply and the signal generator.

### Magnetoelectric coupling coefficient measurements

A three-electrode electrochemical cell was used to measure the potential required to maintain the surface charge, which fluctuated according to the electric polarization of the MENDs in the presence of a magnetic field. To fit within the 8 mm gap of the C-shaped electromagnet described above, a nuclear magnetic resonance (NMR) tube (8 mm outer diameter 0.5 mm thickness) was used as the electrochemical cell. Within the cell, the Ag/AgCl reference electrode, Pt counter electrode, and working electrode were immersed in Tyrode’s solution. The working electrode was prepared by drop-casting MEND solution onto a 0.3×1.2 cm^2^ indium tin oxide (ITO) glass substrate and connecting it to a copper wire using conductive silver epoxy (MG Chemicals 842AR-15ML). The electrical connection was then sealed with epoxy (H.B. Fuller, 10010217) for insulation. The thickness of the MEND film on the ITO substrate was measured via a profilometer (Bruker Dektak DXT-A Stylus). All electrochemical measurements were performed using a potentiostat (Gamry Interface 1010E).

### Fluorescent imaging in cultured hippocampal neurons

All animal procedures were approved by the Massachusetts Institute of Technology Committee on Animal Care (MIT CAC, Protocol #2305000529). Hippocampal neurons were extracted from neonatal rat (Sprague Dawley, 001) pups (P1) and dissociated with Papain (Worthington Biochemical). The cells were then seeded on glass slides (5 mm diameter, Bellco Glass 1943-00005) in 24 well plates at a density of 112500 cells/mL. Prior to seeding, the glass slides were cleaned by evaporating ethanol with an alcohol lamp and then coated with Matrigel® (Corning). The cells were maintained in 1 mL Neurobasal media (Invitrogen). Glial inhibition was conducted with 5-fluoro-2’-deoxyuridine (FUDR, F0503 Sigma) 3 days after seeding. 4 days following seeding, the neurons were transduced with 1 µL of an adeno-associated virus serotype 9 (AAV9) carrying a fluorescent calcium ion indicator GCaMP6s under a pan-neuronal human synapsin (hSyn) promoter (AAV9-hSyn::GCaMP6s, Addgene viral prep #100843-AAV9, >1×10^13^ IU/µL). After 5 days of incubation, calcium (Ca^2+^) imaging with GCaMP6s experiments were performed.

For Ca^2+^ imaging, the hippocampal neurons were washed once with Tyrode’s solution and then immersed into 0.1 mg/mL MEND solution within a well of 24-well plate for 1 hour to obtain ∼1 g_MEND_/mm^2^ density of the particles on the neuronal surfaces. To change the MEND density on neuronal surfaces, we varied the concentration and volume of the MEND incubation solution. The hippocampal neurons on the glass coverslips were then transferred into a custom sample holder containing 200 µL Tyrode’s solution, which was introduced into an electromagnet described above. The fluorescence changes in GCaMP6s were recorded at a rate of 1 frame per second (fps). The fluorescence intensity of each cell was analyzed with ImageJ software, and F_0_ values were determined as the average fluorescence intensity during 30s prior to the initial application of the AMF. Averaged ΔF/F_0_ was estimated from 300 randomly selected neurons from 3 plates. The number of the peaks in ΔF/F_0_ was counted when the peak height is larger than the half of mean standard deviation of ΔF/F_0_ for the overall imaging time of each video. The cells were defined as responsive if ΔF/F_0_ value exceeded 3 times the standard deviation (3σ) of the baseline (collected over 30 s prior to the initial application of the AMF) within 15 sec from MF onset.

To examine the cell viability in the presence of MENDs following MF stimuli, we performed analysis with a Live/Dead Viability/Cytotoxicity Kit (Invitrogen, L3224). The live and dead cells were indicated with Calcein AM, green and ethidium homodimer-1, red. The proportion of viable cells calculated by normalizing the viable cell number to the total number of cell nuclei based on Hoechst staining (Thermo Scientific, Hoechst 33342 Solution (20 mM)). The neurons were incubated with the assay reagents for 20 min in 37 ℃ and then imaged before and after three cycles of MF field via a fluorescent microscope (Olympus IX73, 20x objective lens).

### Stereotactic surgeries

All animal procedures were approved by the MIT CAC (Protocol #2208000413). Surgeries were conducted under aseptic conditions within a stereotaxic frame (David Kopf Instruments). Mice were anesthetized under isoflurane (0.5-2.5% in O_2_) using an anesthesia machine (VET EQUIP). During the surgery, the eyes were covered with ophthalmic ointment, and a heat pad was used to maintain the animals’ core temperature. Mice were provided with subcutaneous injections of 0.6 mL of sterile Ringer’s solution for hydration and extended-release buprenorphine (1 mg kg^-1^) for analgesia at the start of the procedure. The head was fixed into position using ear bars, then the fur on the top of the head was removed using depilatory cream. Following sterilization of the skin with betadine and ethanol, a midline incision was made along the scalp. Coordinates for the injection/implantation site were established based on the Mouse Brain Atlas^81^. A small craniotomy was drilled through the skull using a rotary tool (Dremel Micro 8050) and a carbon steel burr (Heisinger, 19007-05), and the dura was gently removed. Particles were injected into the VTA (anterior-posterior, AP –2.9 mm, medial-lateral, ML –0.5 mm, dorsal-ventral, DV –4.5 mm) using a microinjection apparatus (10 µL Hamilton syringe #80308, UMP-3 syringe pump, and Micro4 controller; all from World Precision Instruments).

For c-Fos expression analysis and behavioral assays, 1.5 µL of MEND particles, MND control particles, or PBS were injected at a rate of 500 nl min^−1^. For MENDs and MND nanoparticle solutions, we used 1.5 mg mL^−1^ concentration for all experiments, except for an additional 0.5 mg mL^−1^ low concentration of MENDs tested in the c-Fos expression experiments. For PBS controls, the same volume of sterile phosphate buffered saline was injected instead of particles. After injection, the syringe was lifted up by 0.1 mm from the initial DV coordinate and left in place for 10 min before slowly withdrawing (0.5 mm min^−1^), and then the skin was sutured (Ethicon, Ethilon 661H, polyamide 6, 19 mm needle length).

For the mice used for fiber photometry experiments, 300 nL of AAV9 hSyn::GCaMP6s, Addgene viral prep #100843-AAV9, >1×10^13^ IU/µL) was injected along with the 1.3 µL particles or PBS. The AAV solution was loaded into the syringe tip after the particles or control PBS, such that it was injected first and immediately followed by the particle or PBS injection into the brain. Following the injection, the syringes were lifted as described above. At 0.1 mm above the injection coordinates, a 200 µm-diameter silica fiber (Thorlabs FT200EMT) with a 2.5 mm diameter stainless steel ferrule (Thorlabs SF230-10) was implanted and cemented in place with adhesive acrylic (C&B-Metabond, Parkell) and followed with dental cement (Jet Set-4).

Following surgery, a subcutaneous injection of carprofen (5 mg kg^-1^) was provided as an anti-inflammatory and analgesia agent, and the animals were placed in a clean recovery cage, part of which was positioned on a heating pad with ad libitum access to water and diet gel wet food.

### Fiber photometry

Fiber photometry recordings were performed using a Neurophotometrics fiber photometry system (FP3002, Neurophotometrics LTD). This system utilized a blue (peak wavelength λ = 470 nm) light-emitting device (LED) to excite GCaMP6s and a violet (peak λ = 415 nm) LED as an isosbestic wavelength control, with a fluorescence light path that includes a dichroic mirror to pass emitted green fluorescence (passband 495 – 530 nm) onto a complementary metal-oxide semiconductor camera (FLIR BlackFly). The 470 nm and 415 nm LEDs were each calibrated to provide 75 µW of optical power out of the tip of a 200 µm silica fiber matching the type implanted into the animals (Thorlabs FT200EMT). The system was coupled to a low-autofluorescence branching bundle patch cord (400 µm core, 0.57 NA, Doric), which was connected to the animal’s implant using a ceramic split mating sleeve (Thorlabs ADAF1). All photometry recordings were performed under isoflurane anesthesia (0.5-2.5% in O_2_, VET EQUIP) using a nose cone. After induction, the patch cord was connected to the animal’s fiber implant, and the animal’s head was placed in the center of the custom magnetic apparatus described above: 10 cm diameter solenoid AMF coil with a ring-shaped static magnet inside to provide OMF, whose centers are aligned. Fiber photometry recordings were performed at 130 Hz sampling rate. After recording a 5-minute baseline to account for rapid photobleaching at the start of the experiment, 2s-pulses of magnetic field (OMF 220 mT, AMF 150 Hz, 10 mT) were applied every 180 s. Each animal received 10 total pulses of magnetic stimulation. Data were collected in Bonsai software and exported to MATLAB (MathworksR2022a) for analysis. Photometry data were analyzed as follows: data were low-pass filtered below 25 Hz (2^nd^ order Butterworth filter), both isosbestic (405 nm excitation wavelength) and Ca^2+^-sensitive (470 nm excitation wavelength) signals were fitted with MATLAB exp2 fitting function, and then the fitting functions were subtracted from the original signals to correct for photobleaching. To remove the motion artifacts, the baseline corrected isosbestic signal was subtracted from the baseline-corrected 470 nm signal. The fluorescence average of 30 s prior to the first stimulation epoch was taken as F_0_, and ΔF/F_0_ was segmented corresponding to each stimulation epoch (30 s pre-stimulation and 150 s post-stimulation).

### Immunohistochemical quantification of biomarkers

For the c-Fos expression experiments, mice were anesthetized one week after the surgery via an intraperitoneal injection of ketamine (100 mg/kg) and xylazine (10 mg/kg) mixture. The heads of the anesthetized mice were placed into the center of a custom-built apparatus with a 10 cm diameter solenoid AC coil outside and ring-shaped static magnet inside, whose centers are aligned. Each mouse received three 2s-pulses of magnetic stimulation (220 mT OMF; 150 Hz, 10 mT AMF) separated by 90 min rest epochs prior to sacrificing. Following the 90-min c-fos induction period, the anesthetized mice were euthanized by a lethal intraperitoneal injection of a sodium pentobarbital (Fatal-plus, 50 mg mL^−1^, dose 100 mg/kg). The mice were then transcardially perfused with PBS and 4% paraformaldehyde (PFA), and their brains were extracted and kept in 4% PFA overnight at 4 ℃. After moving the fixed tissue to PBS and storing at 4 ℃ for 24 hours, the brains were sectioned into 50 µm coronal slices with vibrating blade microtome (Leica VT1000S). The permeabilization was done on the slices for 30 min in the dark at room temperature in a 0.3% v/v Triton X-100 solution in PBS, and then the blocking was done for 1 hour with 0.3% v/v Triton X-100 and normal donkey serum 5% v/v blocking serum solution in PBS with an orbital shaker. Following PBS washing three times, the brain slices were incubated in the first antibody solution overnight at 4 ℃. After three washes with PBS, the brain slices were immersed in a secondary antibody solution for 2 hours at room temperature on the orbital shaker in a dark room. After another three washes with PBS, the brain slices were stained with DAPI, washed, and then transferred onto glass slides with mounting medium (Fluoromount G, Southern Biotech). Rabbit anti-cfos (1:500, Cell Signaling Technology, 2250s) primary antibodies and Donkey anti-rabbit Alexa Fluor 488 (1:1000, Invitrogen, A-21206) secondary antibodies were used for the c-fos expression analysis. Three pairs of primary and secondary antibodies were used for the toxicity assessment and summarized in **Table S2**.

**Table S2.** Antibodies used for biocompatibility analysis.

| **Antibody** | **Primary/Secondary (Dilution)** | **Product Information** |
| --- | --- | --- |
| Goat anti-Iba1 antibody | Primary Antibody (1:500) | Abcam ab107159 |
| Goat anti-GFAP antibody | Primary Antibody (1:1000) | Abcam ab53554 |
| Rabbit anti-CD68 antibody | Primary Antibody (1:250) | Abcam ab125212 |
| Donkey anti Rabbit IgG (H+L), Alexa Fluor 488 | Secondary Antibody (1:1000) | Thermo Scientific A-21206 |
| Donkey anti Goat IgG (H+L), Alexa Fluor 633 | Secondary Antibody (1:1000) | Fisher Scientific A-21082 |

### Behavioral assays

For behavioral experiments, the mice were tested during the light phase of the 12-hour light/dark cycle. For the place preference assay, 71 mm inner diameter acrylic cylinder was divided into three chambers – two 11 cm long stimulation chambers (solid and stripe-patterned bottom) separated by a 19 cm long neutral chamber (purple colored bottom). The stimulation chambers were equipped with custom-made electromagnets and static magnets outside the stimulation chambers, as shown in **Fig. 4j**, Fig. S26. For three days (Day –2 to Day 0, **Fig. 4j**, Fig. S26) before the baseline place preference test, the mice were habituated daily for 15 min in the setup and to the researchers. On Day 1, the mice were placed into the chamber for 5-min habituation during which they could explore the chambers freely without any stimulation, and their place preference was tested for the following 10 min without any magnetic stimulation. From Day 2 to Day 4, following a 5-min habituation in the setup, the mice were stimulated with the magnetic field when they entered the less-preferred area for 10 min daily. On Day 5, the place preference was tested again without any stimulation. The mouse location in the arena was recorded with two cameras (Logitech HD Pro Webcam C920) positioned at the ends of each stimulation chamber and Logitech Capture Bio Recording & Streaming Software (2.08.11) was used to record the videos. Analysis of the place preference assay was done manually, where the observer was blinded to the subject type and marked the time spent in each chamber. Mice that showed more than 500 s (total test time is 600s) preference to a chamber during pre-test or stayed 0 s at the stimulated chamber on Day 2 were eliminated from the subsequent analyses.

### Statistical analyses

OriginPro 2019 was used for assessing the statistical significance of all comparisons in this study. Although sample sizes were not determined with the power analysis, the group sizes for immunohistochemistry and behavior tests were decided to be similar with previous research performed in the same brain circuit. For c-Fos expression analysis, the data distribution was tested for normality, and then analyzed with ANOVA followed by Tukey’s post-hoc comparison test (*P < 0.05, **P < 0.01, ***P < 0.001). For the immunohistochemistry analyses relevant to the toxicity assessment, unpaired t-test was used to assess the differences between two groups, where significance threshold was indicated with n.s., P > 0.05, *P < 0.05, **P < 0.01, ***P < 0.001. For behavioral experiments, a Kruskal Wallis followed by Tukey’s post-hoc comparison test was applied to compare three groups simultaneously with thresholds of n.s. P > 0.05, *P < 0.05, **P < 0.01, ***P < 0.001, ****P < 0.0001. For the comparison of Pre– and Post-learning days within each group, paired t-test was performed when the data distribution was found normal. For non-normal distribution, Wilcoxon Signed Racks Test was performed. For the preference change data comparing all three groups, the data followed normality, and thus one-way ANOVA was used followed by Tukey’s post-hoc comparison test.

