## Supplementary Information for "Magnetoelectric Nanodiscs Enable Wireless Transgene-Free Neuromodulation"

**Supplementary Note 1**

The magnetoelectric coefficient, $\alpha_{ME}$, of the core-double shell $Fe_3O_4$-$CoFe_2O_4$-$BaTiO_3$ magnetoelectric nanodiscs (MENDs) was measured to be 150 mV $mT^{-1}$ $cm^{-1}$ at the combined magnetic field (MF) conditions: offset magnetic field (OMF) 220 mT and alternating magnetic field (AMF) with a frequency $f_{AC}$=150 Hz and amplitude 10 mT (**Fig. 1h-j**). Using the particle diameter of 231 nm and the AMF magnitude of 10 mT, the potential generated by an individual MEND at these MF conditions can be calculated as ~35 μV ($\frac{150\, mV}{mT\cdot cm} \times 200\, nm\ \times 10\, mT = 34.7\, \mu V$), which is significantly below the threshold for neuronal activation and action potential firing (15-30 mV). To investigate the mechanism underlying the MEND-mediated neuronal modulation we developed a model based on: (1) The instantaneous change in membrane potential, $\Delta V$, for each pulse of the AMF resulting from the integration of voltages generated by individual MENDs distributed with a spacing $d$ on the cell membrane and (2) the dynamic change in membrane potential $V(t)$ resulting from the temporal summation of $\Delta V$ across subsequent periods of the AMF.

To model our system, we leverage the solutions to the three-dimensional cable equation, where neuron processes are approximated as cylindrical bodies, with radius $a$[2]. This framework enables the calculation of the change in membrane potential in the presence of multiple point sources of current or voltage (e.g. microelectrodes, or in our case MENDs) distributed across the membrane surface. As the classic cable model is limited to two microelectrodes, we have expanded it to include a larger number of sources for the membrane potential fluctuations. This allowed us to calculate the change in membrane potential $\Delta V$ as a function of the distance between MENDs, $d$, and the potential generated by a single MEND, $V_0$.

We modeled a neuron receiving stimulation via MENDs on its membrane as a cylindrical cell receiving current injection from a series of microelectrodes with spacing $d$ and zero circumferential angle ($\theta = 0$) aligned along the cell surface (**Fig. S16**). For two microelectrodes separated by the distance $d$ that supply current $i_0$, the change in membrane potential, $\Delta V$, according to the steady state solution of the cable model is:

$$\Delta V(d, i_0) = 0.5 r_i i_0 a\ (L(d) + S(d)) \qquad \textit{Eq.1}$$

Where $r_i$ is the resistance of a unit length of the interior of the cell against longitudinal current flow, $a$ is the radius of the cell, and $L$ and $S$ represent the spatial decay of potential caused, respectively, by the one-dimensional and three-dimensional spread of current[2]. $L(d)$ is defined as:

$$L(d) = \frac{\lambda}{a} e^{-d/\lambda} \qquad \textit{Eq.2}$$

Where $\lambda$ is the neuron length constant, defined as the length over which the membrane potential decays to $V_0/e$ . Values of $S(d)$ have been empirically determined, and we use values from the literature tabulated for varying values of $d$, shown in Table S1[3].

MENDs on the neuron membrane can be represented as multiple microelectrodes spaced at a distance $d$, which we assume to be much smaller than the overall length of the neuron, $l$. Thus, the change in membrane potential during one stimulation pulse integrated across all microelectrodes becomes:

$$\Delta V(d, i_0) = \left| \int_d^l \frac{dV}{dx} dx \right| = |V(l) - V(d)| \qquad \textit{Eq.3}$$
$$= |0.5 r_i i_0 a \, (L(l) - L(d) + S(l) - S(d))|$$

In the approximation that the potential change at a large distance from the point source approaches zero, we can assume that $L(l) \approx 0$ and $S(l) \approx 0$, which leads to a simplification of Eq. 3 to:

$$\Delta V(d, i_0) \approx |0.5 r_i i_0 a(-L(d) - S(d))| \qquad \textit{Eq.4}$$

Given that MEND-mediated modulation effectively relies on voltage application rather than current injection, we applied a steady-state solution from the cable model to calculate current $i_0$ from the MEND potential $V_0$[4]:

$$i_o = \frac{2V_0}{r_i \lambda} \qquad \textit{Eq.5}$$

where $\lambda$ is the neuron length constant (defined above) determined by $r_i$ and the membrane resistance per unit length. Hence, the integrated membrane potential change for a single pulse of the AMF can be rewritten in terms of $d$ and $V_0$ as:

$$\Delta V(d, V_0) = |0.5 r_i i_0 a(-L(d) - S(d))| = \left| 0.5 r_i \frac{2V_0}{r_i \lambda} a \left[ -\frac{\lambda}{a} e^{-\frac{d}{\lambda}} - S(d) \right] \right| \qquad \textit{Eq.6}$$

The values for $a$, $\lambda$, $r_i$, and $S(d)$, taken from the literature, are tabulated in Table S2[3]. This equation implies that the total change in membrane potential $\Delta V(d, V_0)$ increases with decreasing spacing between the individual particles, $d$, and with increasing voltage generated from individual particles, $V_0$.

Using these insights about membrane potential change stemming from the integrated inputs from spatially distributed MENDs, we develop a framework to determine how the membrane potential varies in time across multiple cycles of AMF. Application of AMF to a neuron decorated with MENDs is anticipated to yield temporal summation of the subthreshold potentials generated by the MENDs during each half-period of the AMF. (Note that since the ME effect does not depend on the AMF sign, the frequency of voltage fluctuations in MENDs is $2 \times f_{AC}$, where $f_{AC}$ is the AMF frequency.) This temporal summation of a series of subthreshold potentials is known as neuronal facilitation[5]. To simulate neural facilitation during magnetoelectric stimulation with MENDs, we adopt a stochastic model of the repetitive activity of neurons[3], and we combine this with the three-dimensional cable model of the voltage distribution derived above (Eq. 6).

When a neuron is subjected to a series of potential pulses with a given time interval between them, the membrane potential, $V(t)$, is the sum of two components: $N(t)$, a noise term that accounts for stochastic fluctuations in membrane potential, and $D(t)$, a function describing the reestablishment of resting membrane potential following each applied potential pulse. Since $N(t)$ is a noise term with a mean of 0 mV, on average, $V(t)$ reaches the threshold for neuron spiking when $D(t)$ does so. $D(t)$ is dependent on the state of the membrane potential at the onset of the applied potential pulse, and it is defined as:

$$D(t) = D_F + (D_I - D_F)e^{\frac{-(t-R)}{k}} \qquad Eq.7$$

Where $D_I$ is the membrane potential at the onset of the applied potential, $D_F$ is the membrane potential in the fully-recovered state dependent on the level of excitatory input, $R$ is the absolute refractory period of the neuron spiking (we use $R = 0.7\ s$), and $k$ is the time constant of decay of the after-potential (we fix $k = 9\ ms$ in this simulation). If we consider the sub-threshold voltage delivered by MENDs to be akin to excitatory input to the neuron, then $D_F$ depends on $\Delta V(d, V_0)$, as defined in Eq. 6. $D_I$ varies depending on history, and it is defined as one-half of the difference between –90 mV and the value of $D(t)$ at the onset of the applied potential [3]. Thus, for each period of the applied AMF, $D_F' = D_{F,initial} + \Delta V$ and $D_I' = (D_F' - 90\,mV)\,/\,2$, with these values getting updated for every cycle of the AMF.

According to this model, subthreshold potentials of the MENDs give rise to a potential change $\Delta V(d, V_0)$ for each period of the AMF, which decays according to $D(t)$ (Eq. 7). With an applied AMF of 150 Hz, subsequent voltage impulses delivered by the MENDs summate over time until the critical threshold is crossed and the neuron fires. Using this model, we find that neurons with a resting membrane potential of –70 mV decorated with MENDs and exposed to 150 Hz AMF will be depolarized to the critical threshold for action potential firing, –55 mV, after ≥ 2s, for a single MEND potential $V_0 > 24\,\mu V$ when inter-MEND spacing $d$ is $0.25a$ (**Fig. 3e-g**).

**Table S3.** $S(d)$ at different values of MEND spacing $d$ in fractions of the axonal radius $a$.[3]

| $\boldsymbol{d}$ | $\boldsymbol{S(d)}$ |
|---|---|
| 0.25a | 3.202 |
| 0.5a | 1.212 |
| 0.75a | 0.598 |
| 1a | 0.327 |
| 2a | 0.042 |

**Table S4.** Parameters used to model MEND-mediated neuronal excitation with AMF.

| **Parameter** | **Value** |
|---|---|
| Axial resistivity, $R_i$ | 150 Ωcm [6] |
| Axial resistance, $r_i$ | $R_i/\pi a^2$ |
| Axonal radius, $a$ | 2 µm [7] |
| Length constant, $\lambda$ | 258 µm [8,9] |

### Supplementary Figures

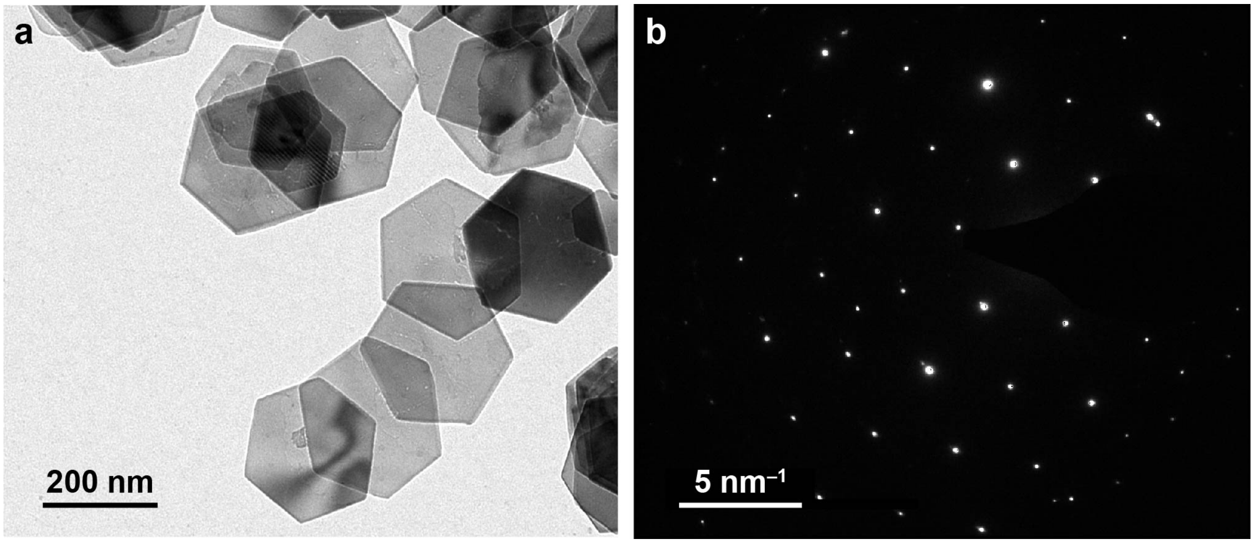


**Fig. S1 | Image of hematite nanodiscs. a,** Transmission electron microscopy (TEM) image of an ensemble of hematite ($Fe_2O_3$) nanodiscs produced via a hydrothermal process prior to reduction. **b,** An electron diffraction pattern of an area on a single hematite nanodisc.

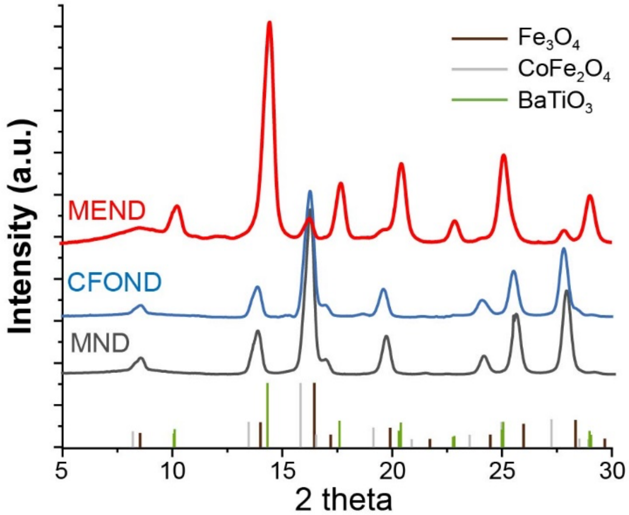


**Fig. S2| X-ray diffraction analysis.** X-ray diffraction spectra of $Fe_3O_4$ magnetic nanodiscs (MNDs, grey), $Fe_3O_4$-$CoFe_2O_4$ core-shell nanodiscs (CFONDs, blue), and co-double shell magnetoelectric $Fe_3O_4$-$CoFe_2O_4$-$BaTiO_3$ nanodiscs (MENDs, red) shown together with the reference spectra for $Fe_3O_4$ (brown, cubic inverse spinel structure), $CoFe_2O_4$ (grey, cubic inverse spinel structure) and $BaTiO_3$ (green, tetragonal crystal system with perovskite structure).

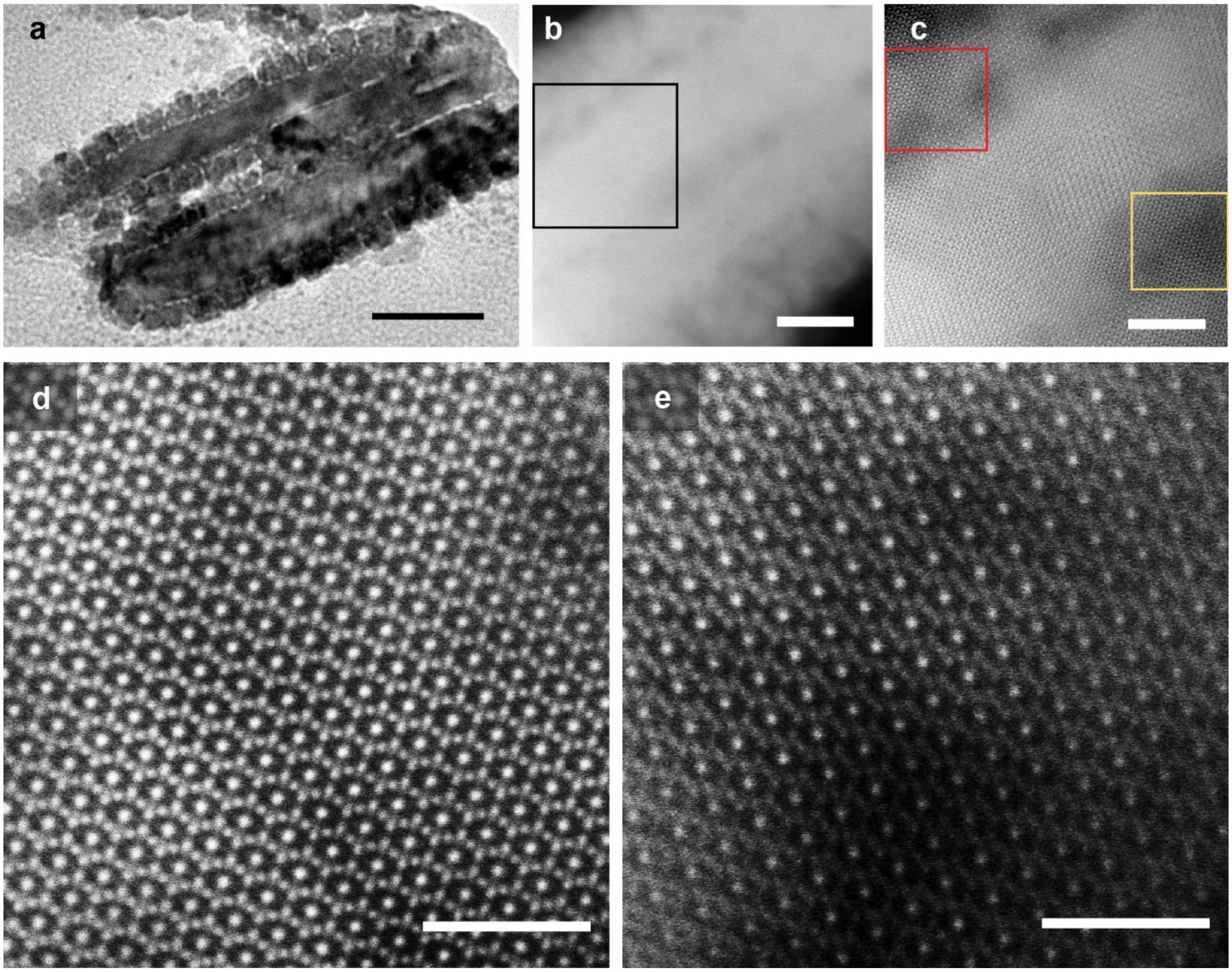


**Fig. S3| Images of cross-sectioned CFONDs. a,** A cross-sectional TEM image of a core-shell $Fe_3O_4$-$CoFe_2O_4$ nanodisc (CFOND). Scale bar = 50 nm. **b-e,** Cross-sectional scanning TEM (STEM) images of a CFOND at different magnifications demonstrate the epitaxial interface between $Fe_3O_4$ and $CoFe_2O_4$ layers. **d,e,** Higher magnification images of the areas marked with red and yellow rectangles in **c**. Scale bars are 10 nm (**b**), 5 nm (**c**), and 2 nm (**d** and **e**).

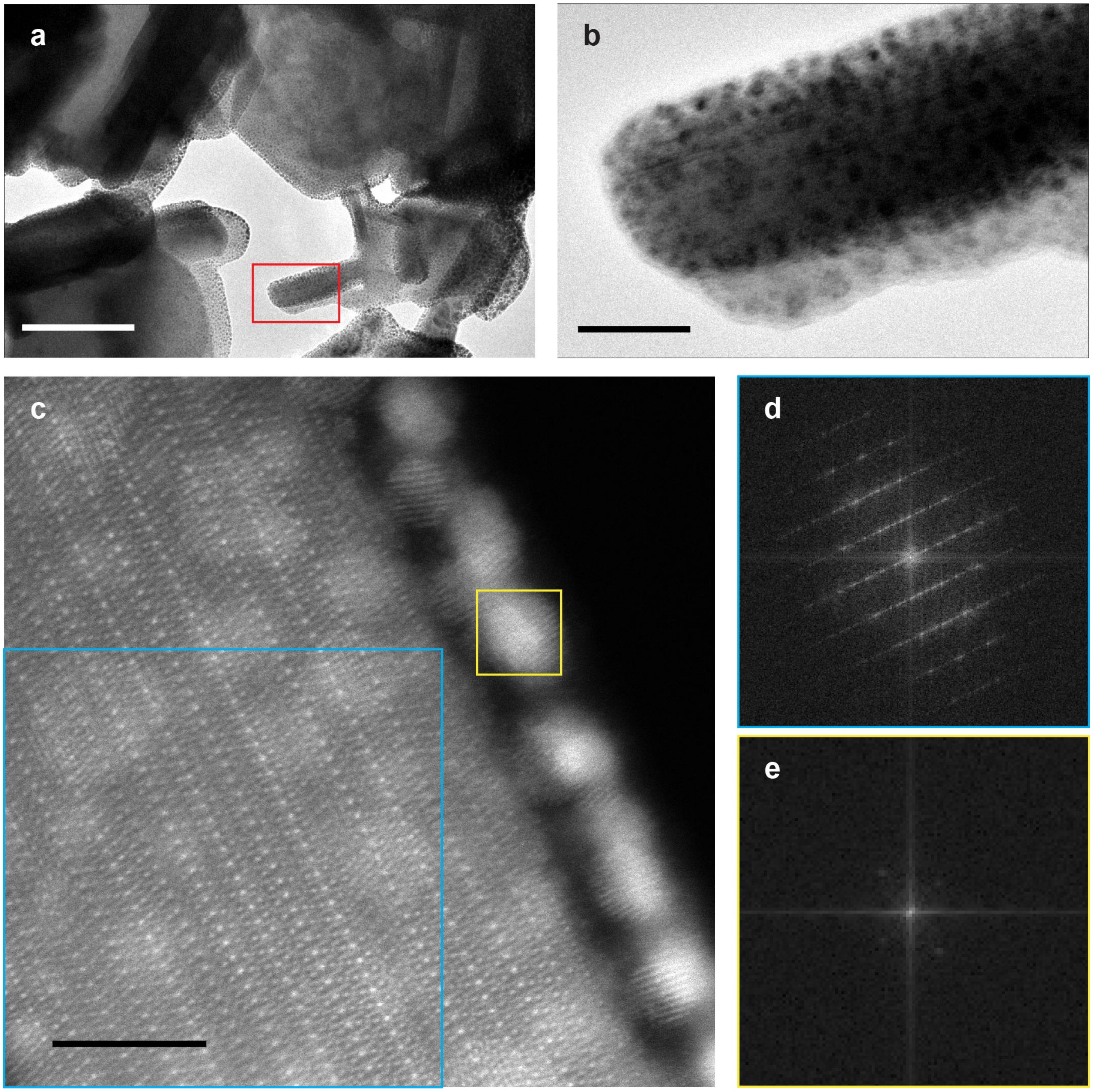


**Fig. S4| Cross-sectioned images of MENDs. a,** Cross-sectional TEM images of a core-double shell $Fe_3O_4$-$CoFe_2O_4$-$BaTiO_3$ magnetoelectric nanodisc (MEND). **b,** A magnified view of the area marked by a red rectangle in (A). **c,** STEM image demonstrates the crystal structure of $BaTiO_3$, (yellow rectangle) which is distinct from that of $Fe_3O_4$ or $CoFe_2O_4$ (inverse spinel, cyan rectangle). **d,e,** Fast Fourier transform images of $Fe_2O_4$-$CoFe_2O_4$ layer (**d**, cyan rectangle in **c**) and $BaTiO_3$ layer (**e**, yellow rectangle in **c**) of the STEM image correspond in **c**. Scale bars are 100 nm (**a**), 20 nm (**b**), 5 nm (**c**).

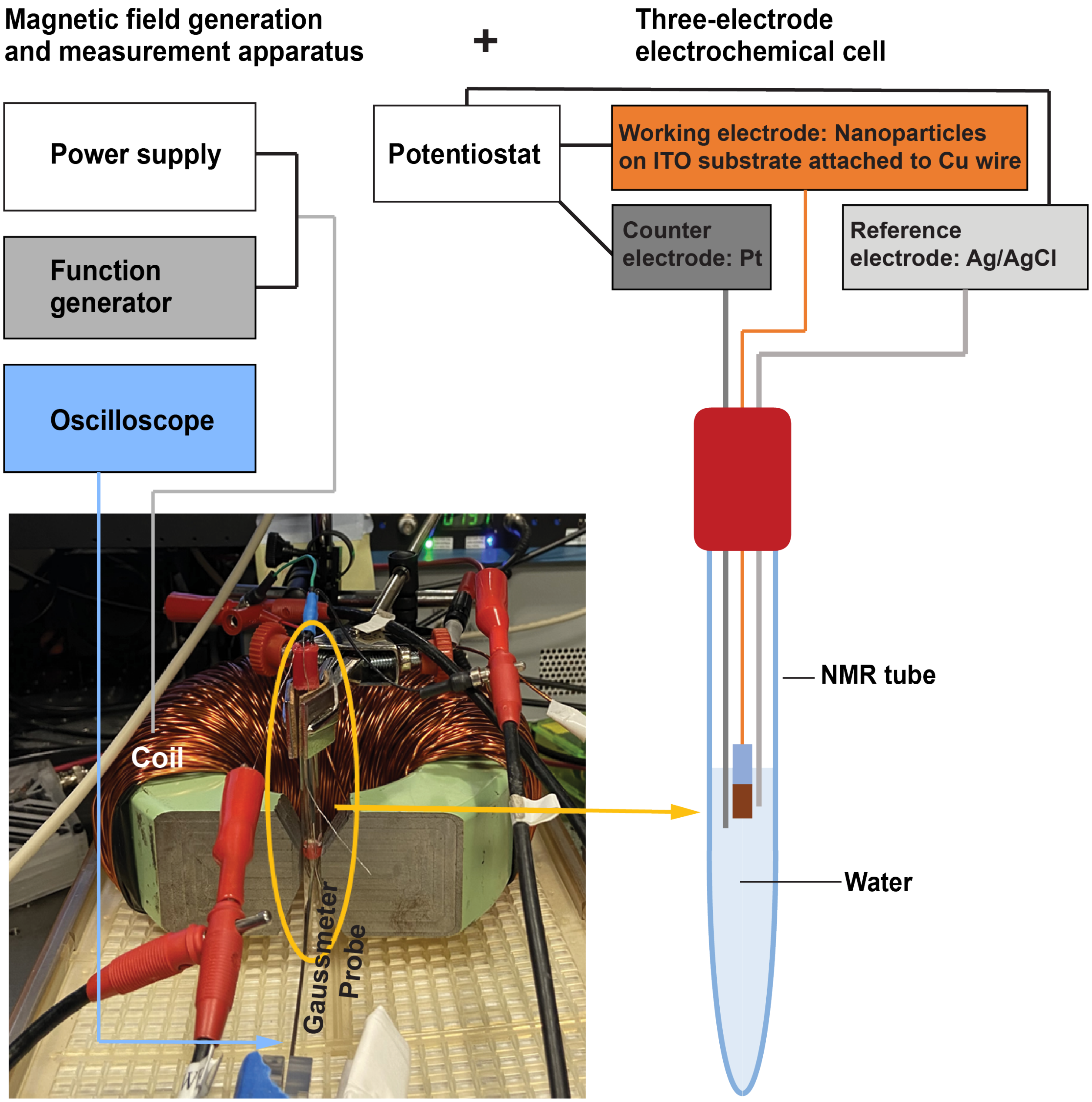


**Fig. S5| Magnetoelectric coefficient ($\alpha_{ME}$) measurement apparatus.** To generate the magnetic field, TEMCo 14 AWG copper magnet wire was wound around a horseshoe-shaped magnetic core with a 0.5-inch gap. The coil was connected to a power supply (CROWN DC-300A Series II) and signal generator (PICOSCOPE 2204A) to generate a compound magnetic field combining an offset magnetic field (OMF, 0 – 320 mT) with an alternating magnetic field (AMF, 0 – 1 kHz, 0 – 14 mT). A nuclear magnetic resonance (NMR) glass tube (8 mm diameter) was used as the electrochemical cell, which was placed into the 8 mm gap of the horseshoe coil. In the cell, the Ag/AgCl reference electrode, Pt counter electrode, and the working electrode were immersed in the Tyrode solution.

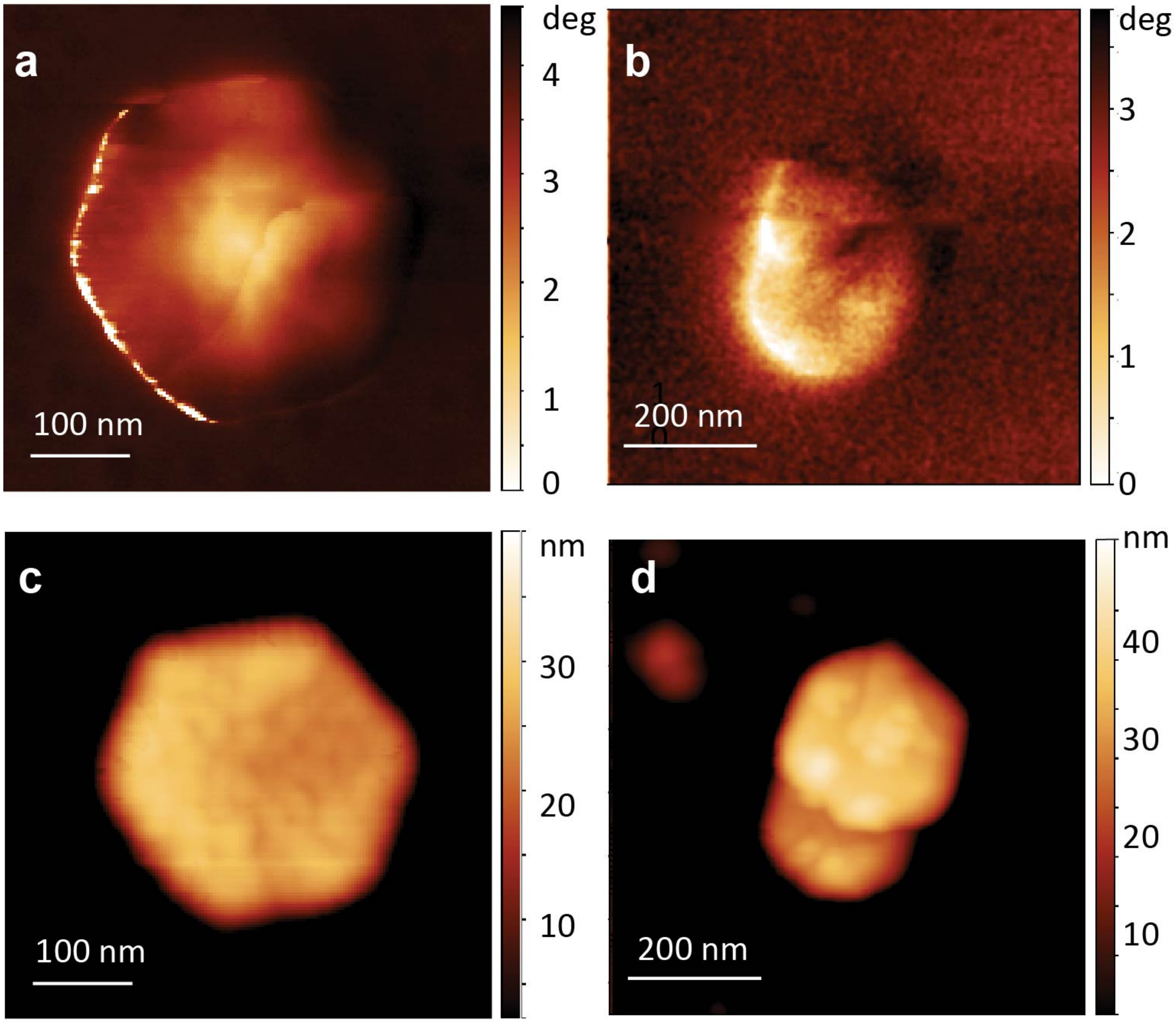


**Fig. S6| Remnant magnetization configuration of CFONDs. a, b,** Magnetic force microscopy (MFM) images of isolated **(a)** and overlapping **(b)** CFONDs. Isolated CFONDs assume the magnetic vortex ground state magnetization, while overlapping CFONDs lose the vortex state in favor of in-plane magnetization. **c, d,** Atomic force microscopy (AFM) images of isolated **(c)** and overlapping CFONDs **(d)** corresponding to the MFM images in (a) and (b), respectively.

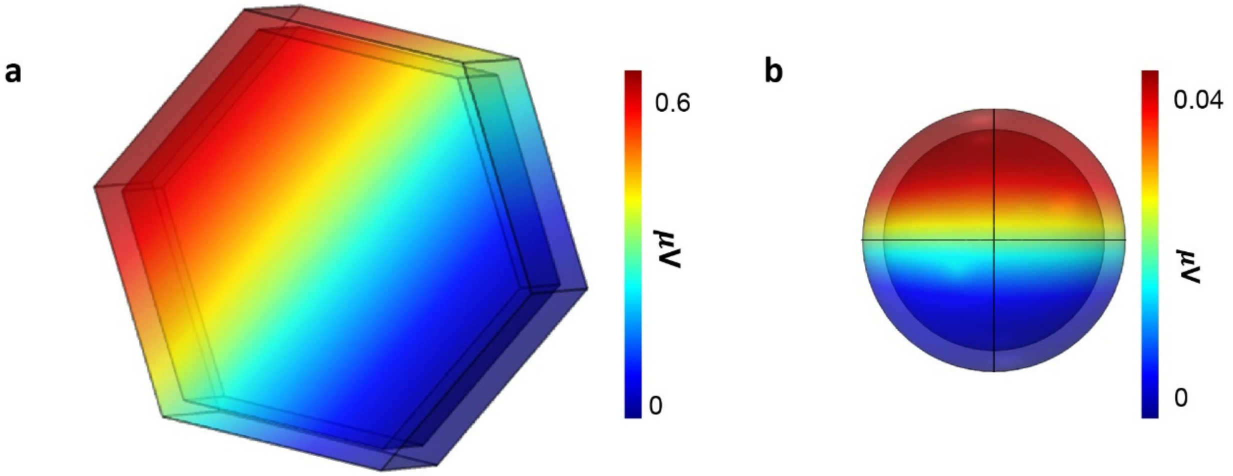


**Fig. S7| Finite element simulation (COMSOL Multiphysics) of the electric polarization in the piezoelectric $BaTiO_3$ shell based on the implementation of the RMS deformation shown in Figure 1f to the saturation magnetization parameter of the core. a,** Hexagonal nanodisc consisting of a 100% $Fe_3O_4$ magnetostrictive core and a 5 nm piezoelectric $BaTiO_3$ shell. **b,** Isotropic nanoparticle consisting of a magnetostrictive core with a volumetric ratio 92% $Fe_3O_4$ and 8% $CoFe_2O_4$ and a 5 nm piezoelectric $BaTiO_3$ shell. The elasticity matrix, coupling matrix, and relative permittivity of $BaTiO_3$ shell has been adapted from prior work.[1]

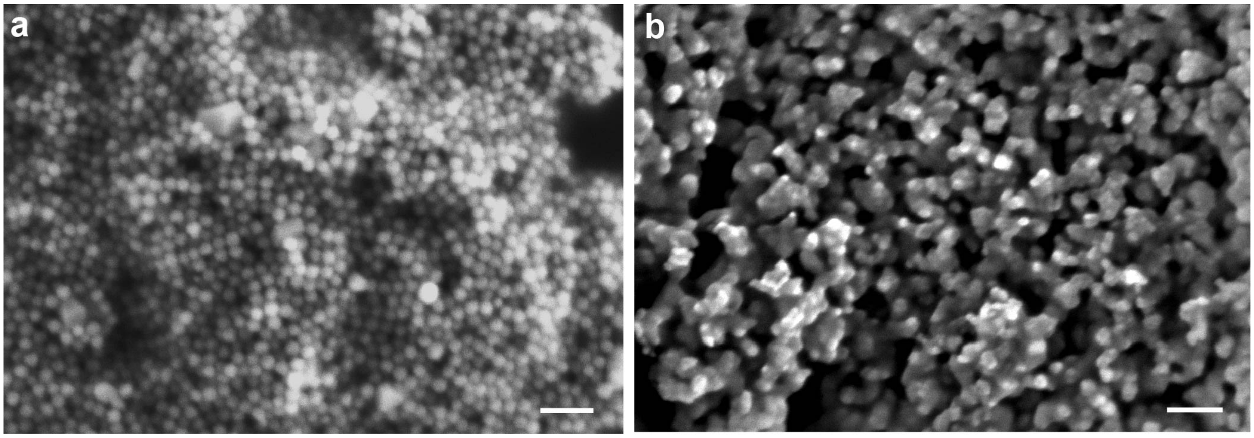


**Fig. S8| Images of isotropically-shaped nanoparticles.** Scanning electron microscopy (SEM) images of isotropic $CoFe_2O_4$ nanoparticles with an average diameter of 25 nm **a,** and core-shell $CoFe_2O_4$-$BaTiO_3$ nanoparticles **b,** produced from cores in **a**. Scale bars = 100 nm.

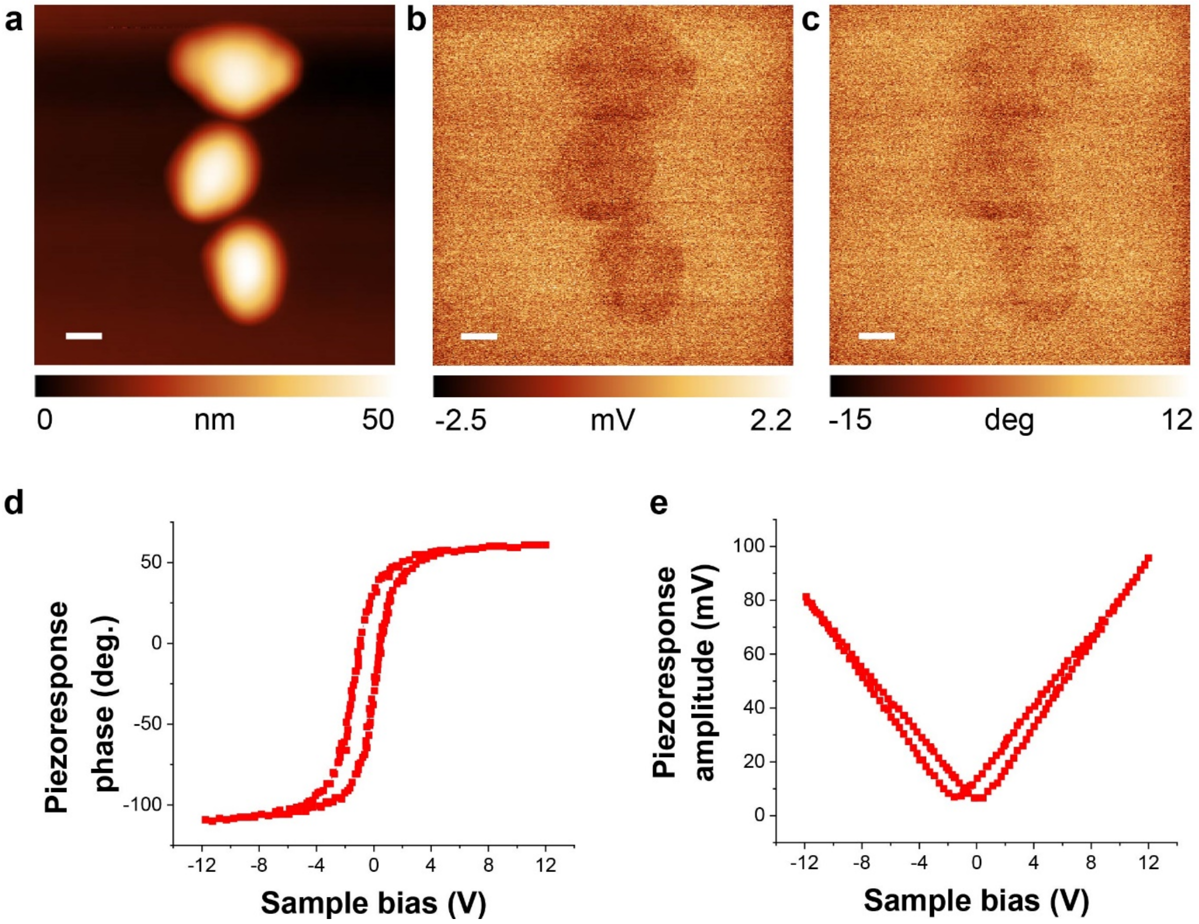


**Fig. S9| Piezoresponse of MENDs.** The images of (**a)** topography, **(b)** piezoresponse amplitude, and **(c)** phase of MENDs observed via piezoresponse force microscopy. **d, e** Piezoresponse phase **(d)** and amplitude **(e)** curves.

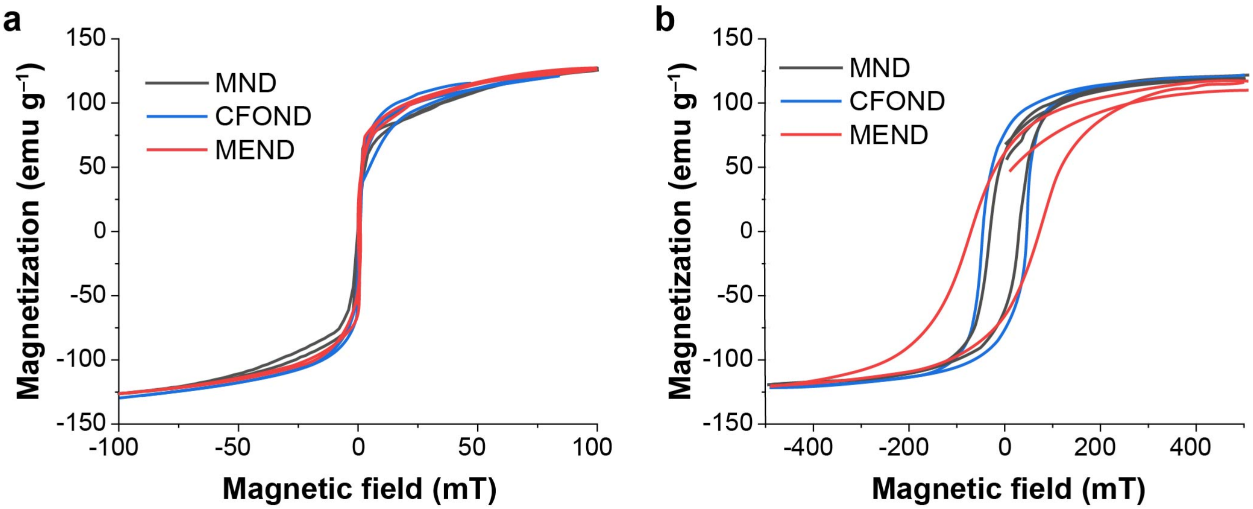


**Fig. S10| Vibrating sample magnetometry (VSM).** VSM of MNDs, CFONDs, and MENDs dispersed in water **(a)** and in a dried pellet form **(b)**.

**Table S5.** Coercivity ($H_C$) and saturation magnetization ($M_S$) of MND, CFOND, and MEND

| | **$H_C$ (mT)** | **$M_S$ (emu/g)** |
|---|---|---|
| **MND** | 38.97±0.76 | 123.34±8.68 |
| **CFOND** | 37.27±4.70 | 113.42±8.97 |
| **MEND** | 66.00±7.37 | 116.64±4.78 |

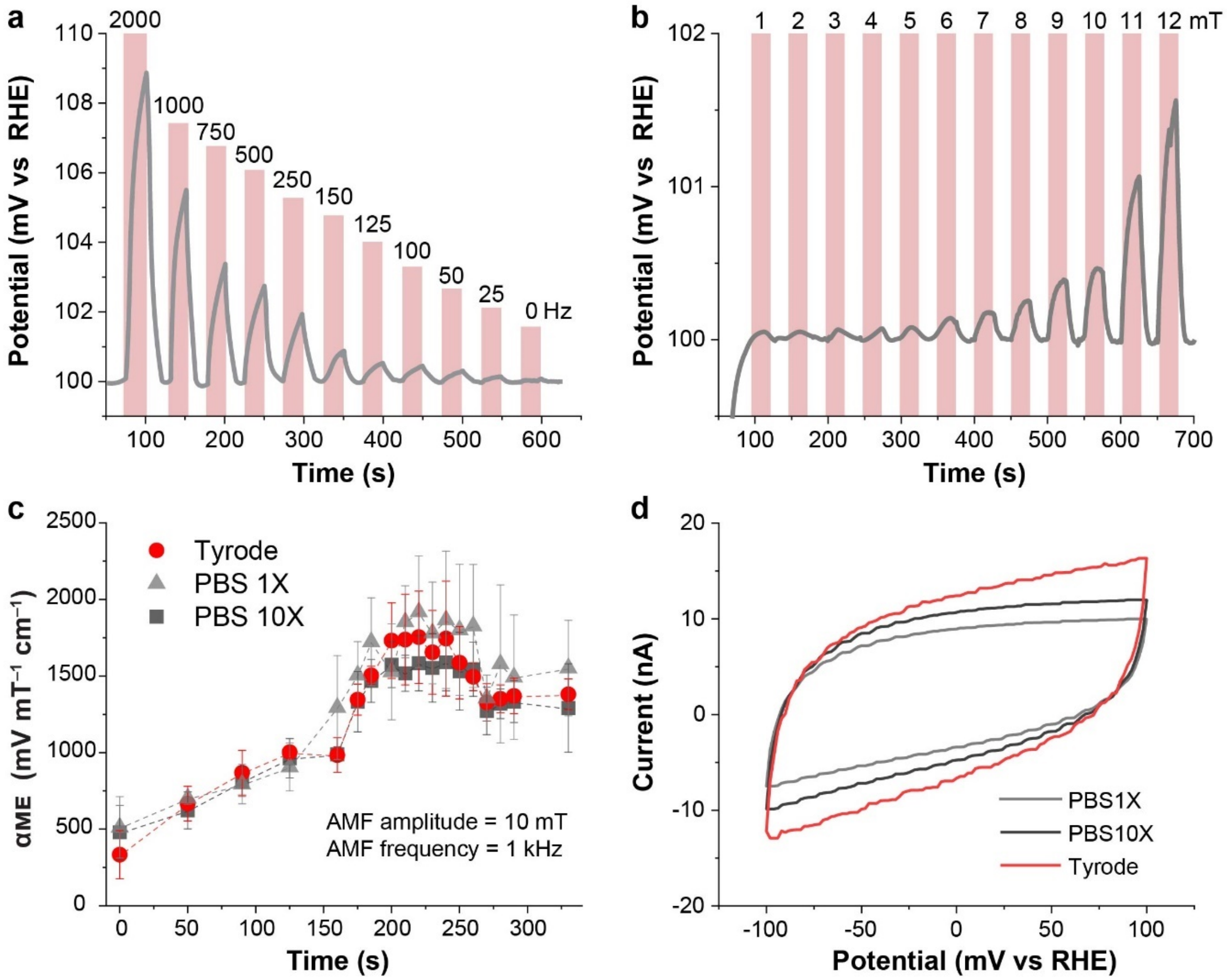


**Fig. S11| Measurement of magnetoelectric coefficients. a,** Changes in potential corresponding to pulses of AC magnetic field with an amplitude of 10 mT and frequencies 25, 50, 75, 100, 125, 150, 250, 500, 1000, and 2000 Hz. **b,** Changes in potential corresponding to pulses of AC magnetic field with a frequency of 150 Hz and amplitudes 1, 2, 3, 4, 5, 6, 7, 8, 9, 10, 11, and 12 mT. Multiple repetitions (n=3) of these measurements were conducted to generate data displayed in **Fig. 1i, j**. The pink boxes indicate the time points when the magnetic field was applied. **c,** $\alpha_{ME}$ at an AMF with a frequency $f_{\mathrm{AMF}}$ =1 kHz and amplitude $H_{\mathrm{AMF}}$ = 10 mT measured at varying magnitudes of OMF for MENDs in Tyrode (red), PBS 1X (light grey), and PBS 10X (dark grey). **d,** The non-faradaic current in cyclic voltammetry (CV) at 25 mV/s scan rate, which implies the electric double layer capacitance within different electrolytes from the equation I = dQ/dt = d($C_{dl}$-E)/dt = $C_{dl}\nu$, where I is non-faradaic current, Q is charge, t is time, $C_{dl}$ is electric-double layer capacitance, E is potential, and $\nu$ is the scan rate of CV.

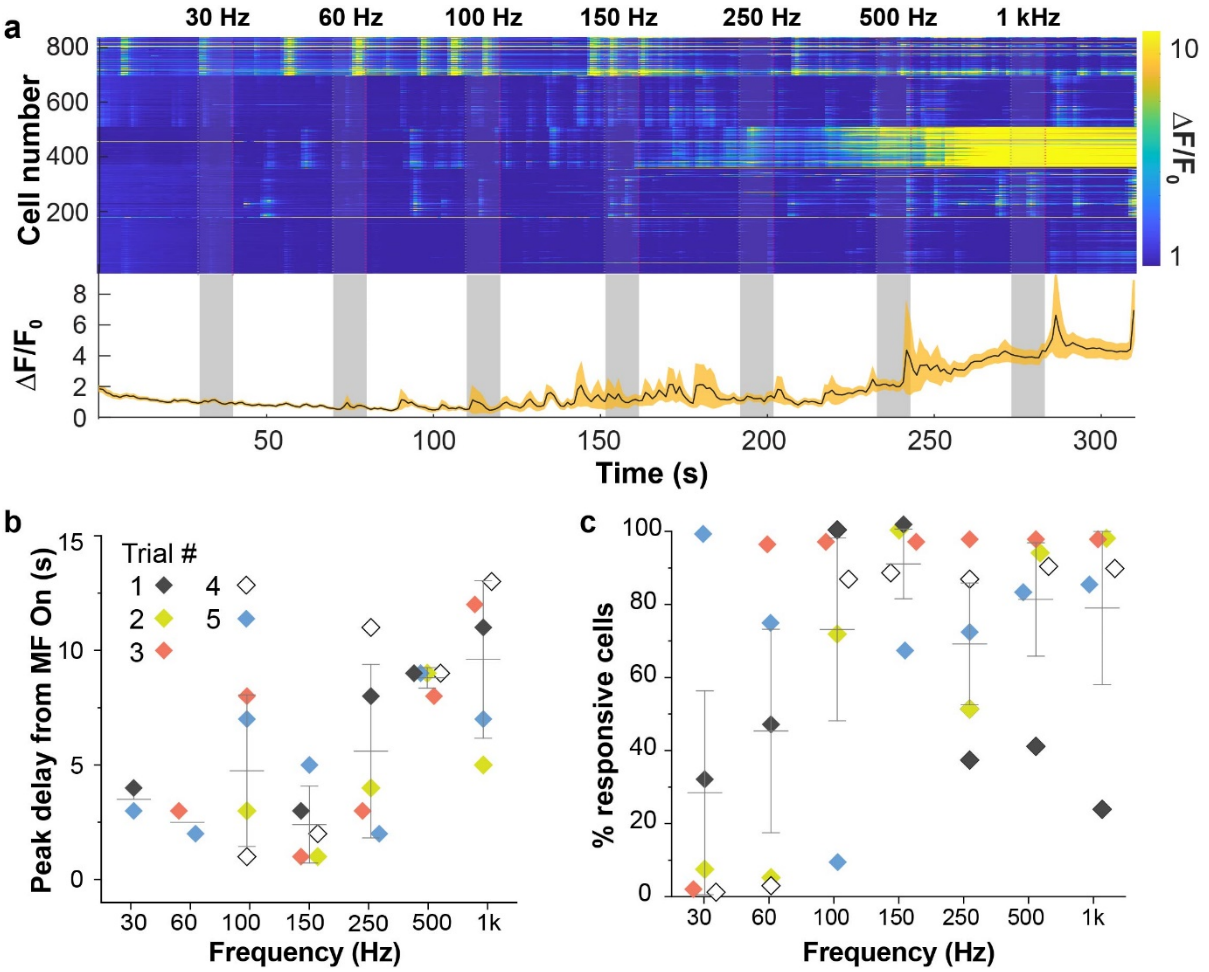


**Fig. S12| GCaMP6s fluorescence imaging at different AMF frequencies. a,** Traces of GCaMP6s $\Delta F/F_0$ of individual (top) and average (bottom) of hippocampal neurons (n=5 culture plates) decorated with MENDs in response to 10 mT AMF with frequencies 30, 60, 100, 150, 250, 500, and 1000 Hz (OMF magnitude 220 mT). The grey bars indicate MF epochs. **b,** Latency of the peak of GCaMP6s fluorescence transient relative to the start of MF epoch. The grey lines represent the average and standard deviation. **c,** The extent of neuronal response to the MF application.

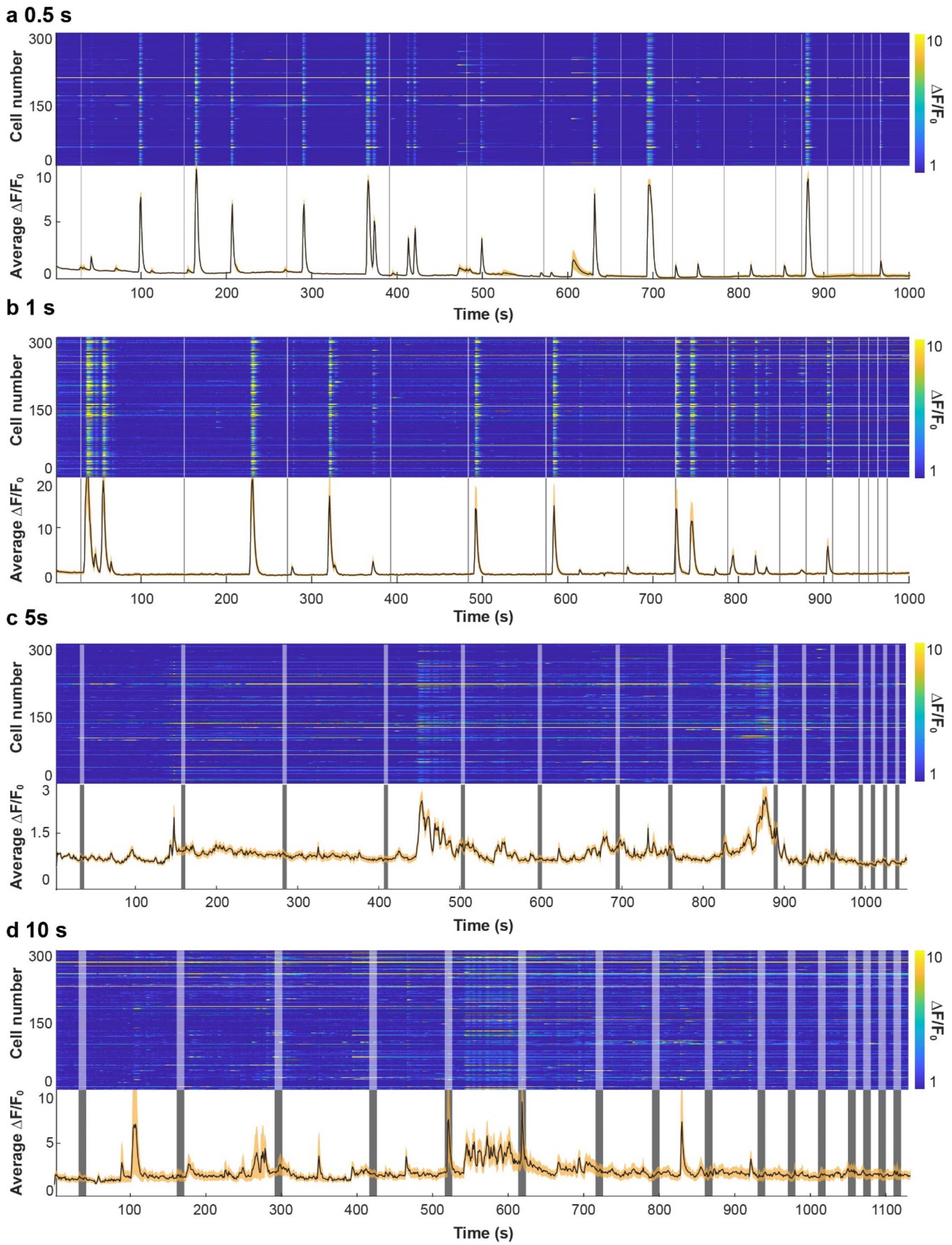


**Fig. S13| Population (top panels, 300 neurons from 3 plates) and average $\Delta F/F_0$ (bottom panels) of GCaMP6s signals for neurons decorated with MENDs and subjected to magnetic field epochs of different length a,** 0.5 s; **b,** 1 s; **c,** 5 s; **d,** 10 s) at intervals of 120 s, 90 s, 60 s, 30 s, 10 s). Grey vertical boxes mark magnetic field epochs (220 mT OMF; 150 Hz, 10 mT AMF).

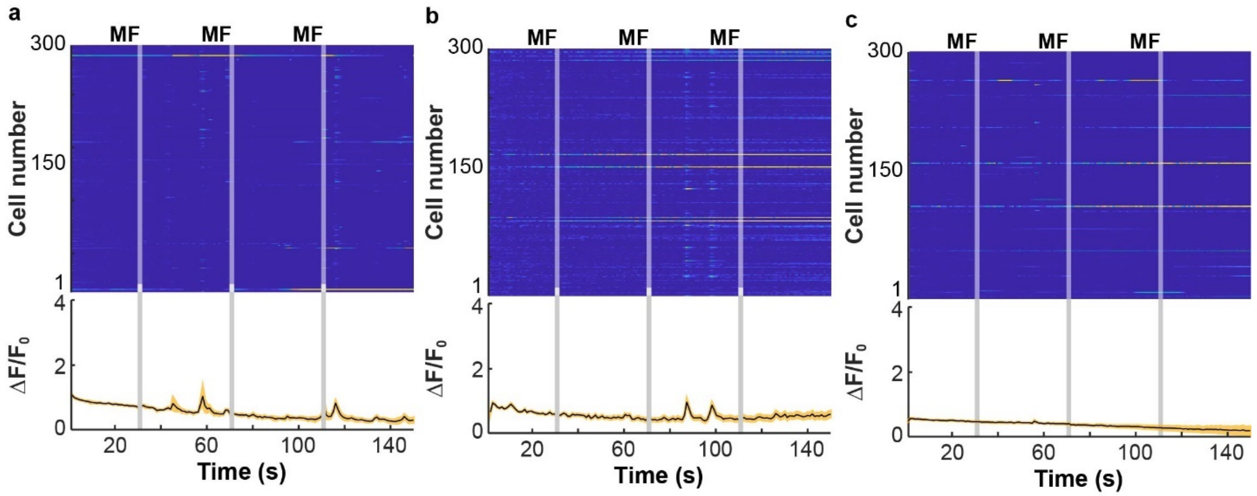


**Fig. S14| GCaMP6s calcium ($Ca^{2+}$) imaging of 300 primary hippocampal neurons (n=3 culture plates).** With MEND densities **a,** 0.5 µg/mm$^2$ **b,** 0.25µg/mm$^2$ **c,** 0µg/mm$^2$. Each row in the top panel represents a neuron, the color bar marks fluorescence intensity change ΔF normalized to baseline fluorescence $F_0$ (averaged between 10-30s). The bottom panels show the average $\Delta F/F_0$ (black) and standard error (yellow shaded area). Vertical grey boxes mark magnetic field epochs (MF, 2 s, 220 mT DC, 150 Hz and 10 mT AC).

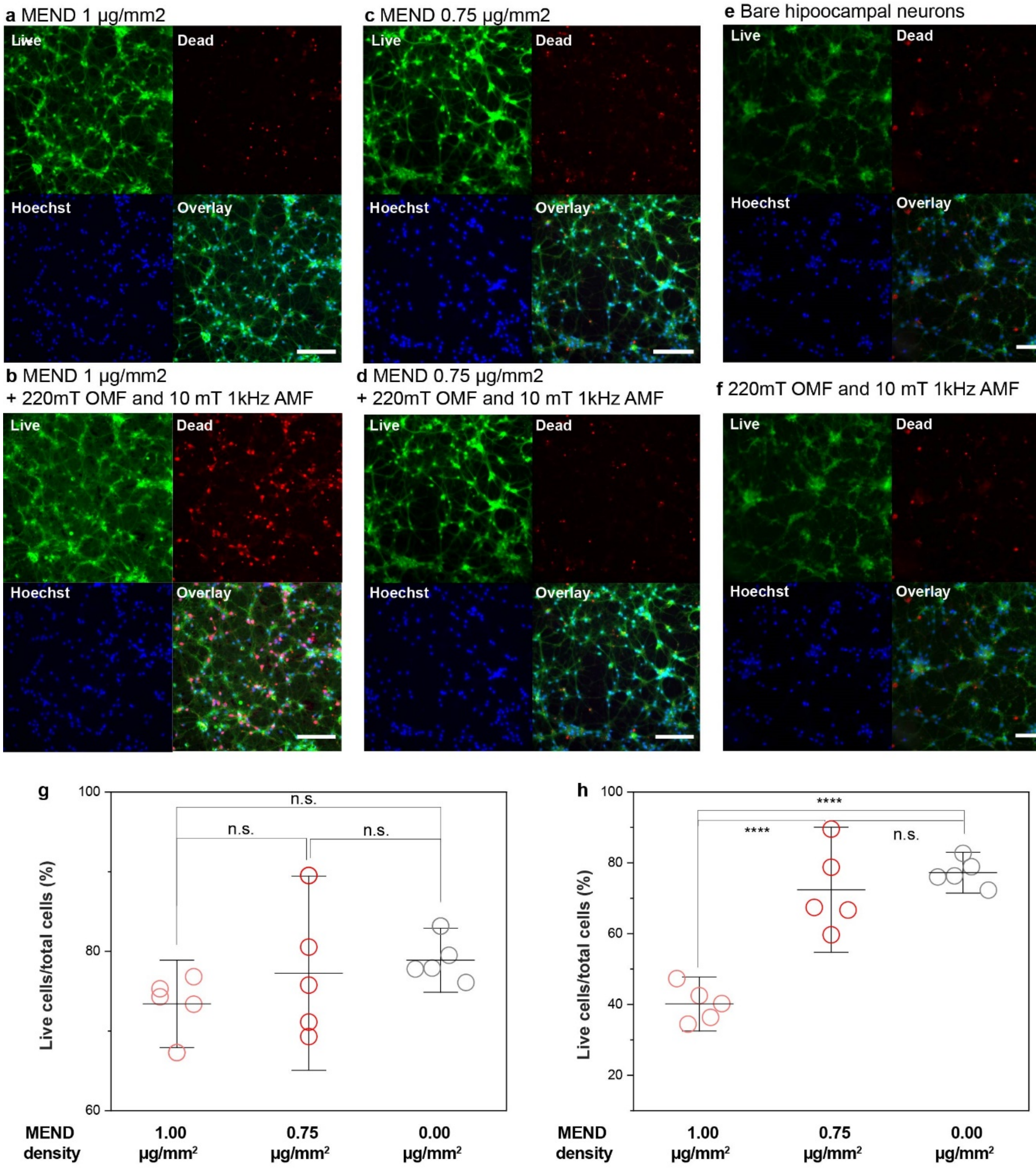


**Fig. S15| Quantification of cell viability.** Live-dead assay of neurons decorated with **a,b,** 1 µg mm$^{-2}$, **c,d,** 0.75 µg mm$^{-2}$, and **e,f,** without MENDs (**a,c,e**) before and (**b,d,f**) after three cycles of MFs. Green – linve cell, Red – dead cells, Blue – nuclei, Scale bar = 150 µm. Live cell numbers normalized with the number of total cells **c,** before and **d,** after MF application on neurons decorated with MENDs at densities of 1 µg mm$^{-2}$, 0.75 µg mm$^{-2}$, and without MENDs. Statistical significance was tested via Kruskal-Wallis ANOVA and Tukey's multiple comparison tests (n = 5 plates per condition, P=0.515 (before) and $6.98\times10^{-5}$ (after) for 1 vs 0.75 µg mm$^{-2}$ ; P=0.278 (before) and $1.78\times10^{-5}$ (after) for 0.75 vs 0 µg mm$^{-2}$ , P=0.881 (before) and 0.598 (after) for 0.75 vs 0 µg mm$^{-2}$ ; ****P≤0.0001, n.s. P>0.05).

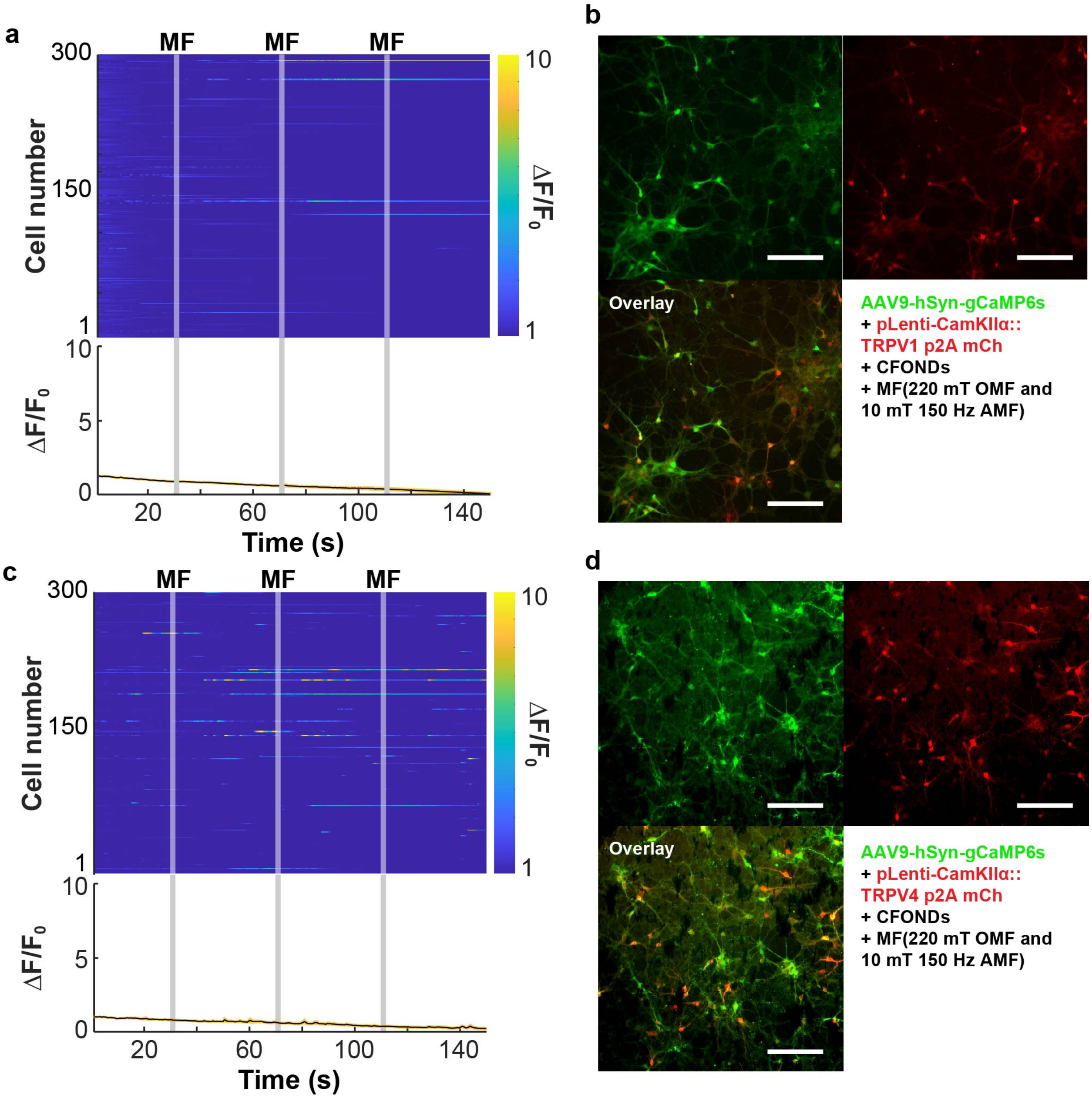


**Fig. S16| GCaMP6s fluorescence imaging of neurons expressing thermo- and mechano-receptors.** GCaMP6s calcium ($Ca^{2+}$) imaging of 300 primary hippocampal neurons (n=3 culture plates) transfected to express **a,** TRPV1 thermoreceptors. and **b,** Image of neurons expressing GCaMP6s (top left), TRPV1 (top right), and their overlay (bottom left). **c,** Ca imaging with neurons transfected to express TRPV4 putative mechanoreceptors. **d,** Image of neurons expressing GCaMP6s (top left), TRPV4 (top right), and their overlay (bottom left). Scale bars are 150 µm.

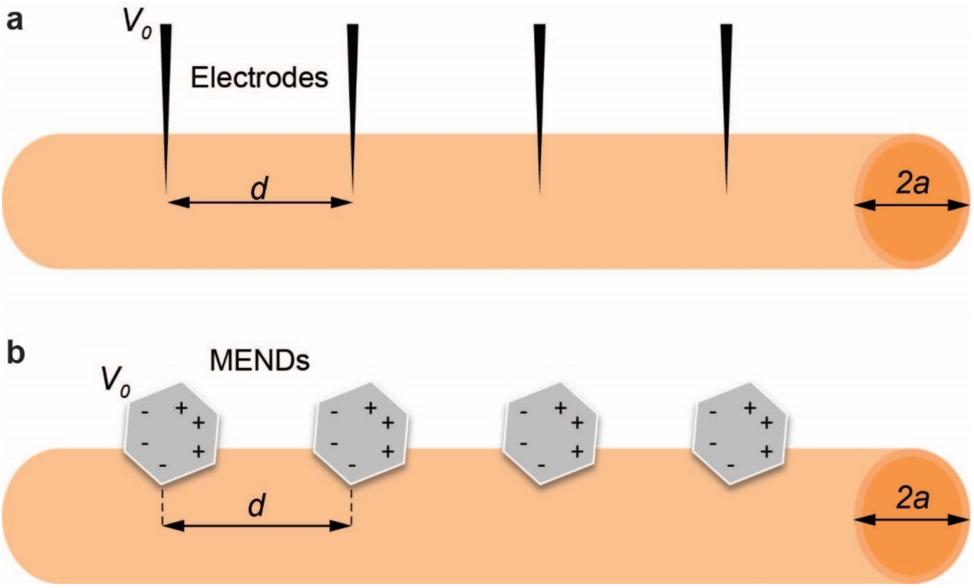


**Fig. S17| Schematic illustration of the model for the mechanistic study. a,** An illustration of the classic cable model in the presence of multiple voltage sources. **b,** An analogous model using MENDs as voltage sources. Here $d$ is the inter-electrode or inter-particle distance, $a$ is the axon radius, and $V_0$ is the potential applied by an electrode or generated by an individual MEND.

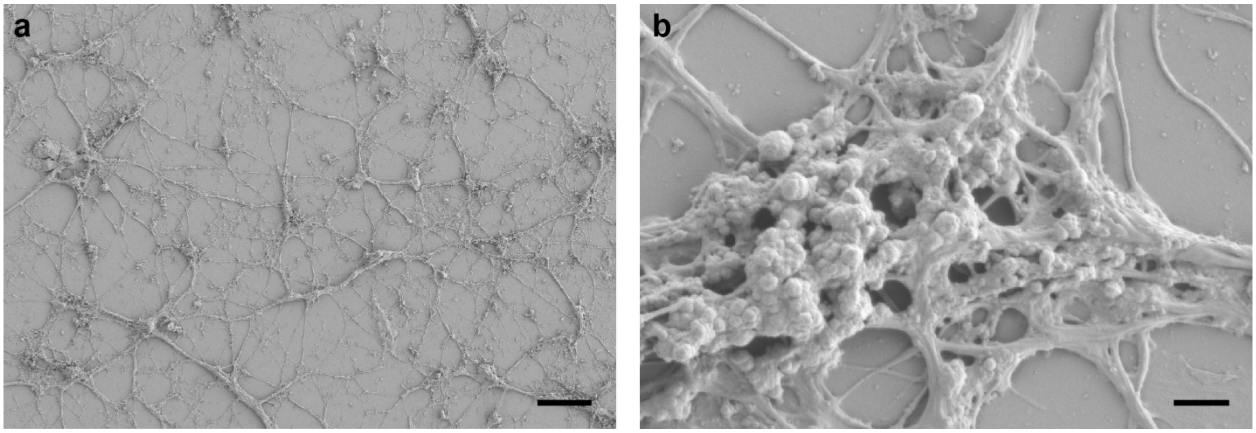


**Fig. S18| SEM images of neurons.** SEM images of primary hippocampal neurons (at 10 days in vitro) without any nanoparticles at **a,** 200X (Scale bar = 50 $\mu$m) and **b,** 4000X magnification. (Scale bars = 2 $\mu$m).

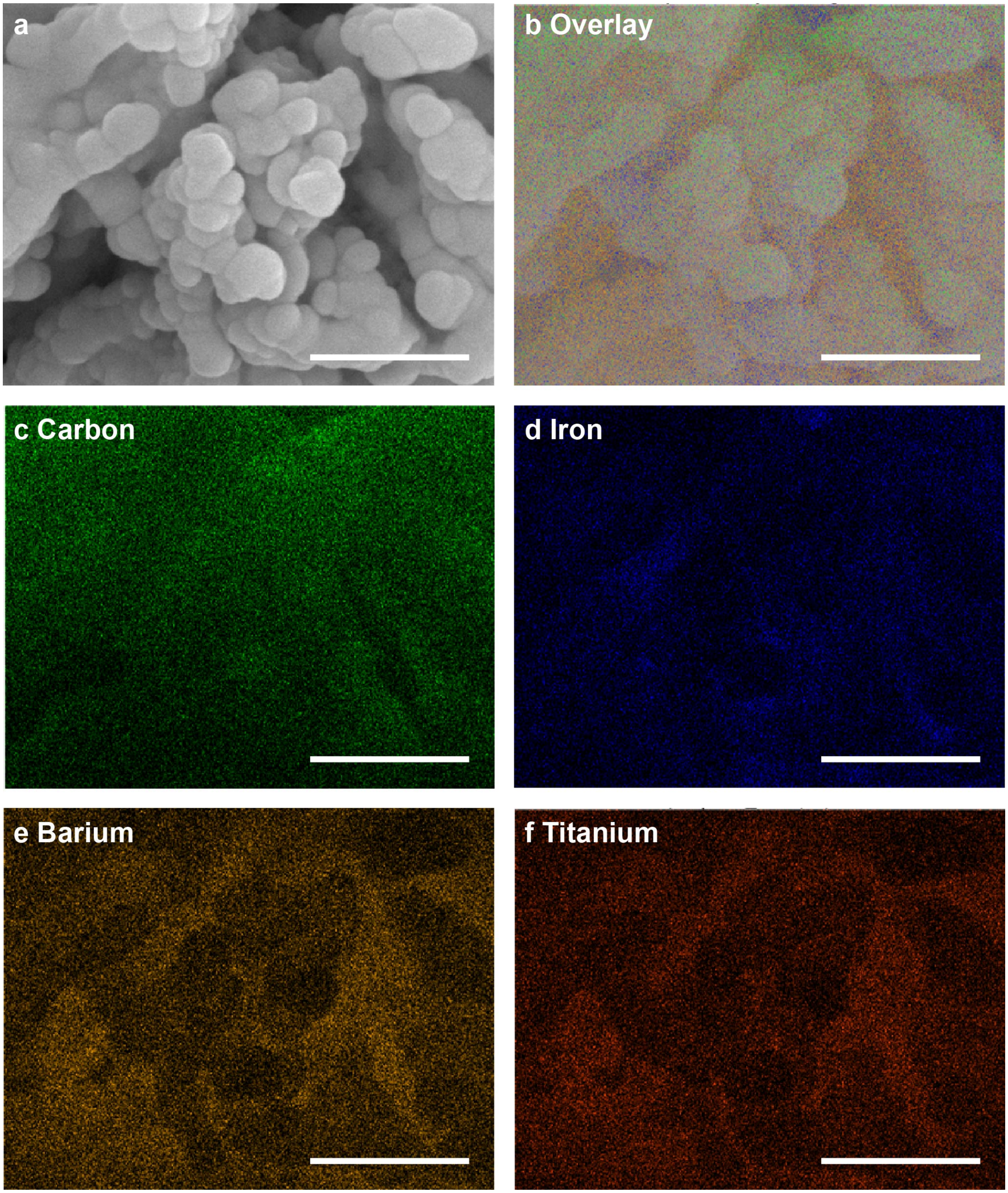


**Fig. S19| Images of neurons decorated with MENDs. a,** SEM image of hippocampal neurons (10 days) decorated with MENDs. **b-f,** Energy-dispersive X-ray spectroscopy mapping on the image in **a**. **b** Merged image of the carbon (C, green, also shown in **c**), iron (Fe, blue, also shown in **d**), Barium (Ba, yellow, also shown in **e**), and titanium (Ti, red, also shown in **f**) atomic maps. Scale bars = 500 nm.

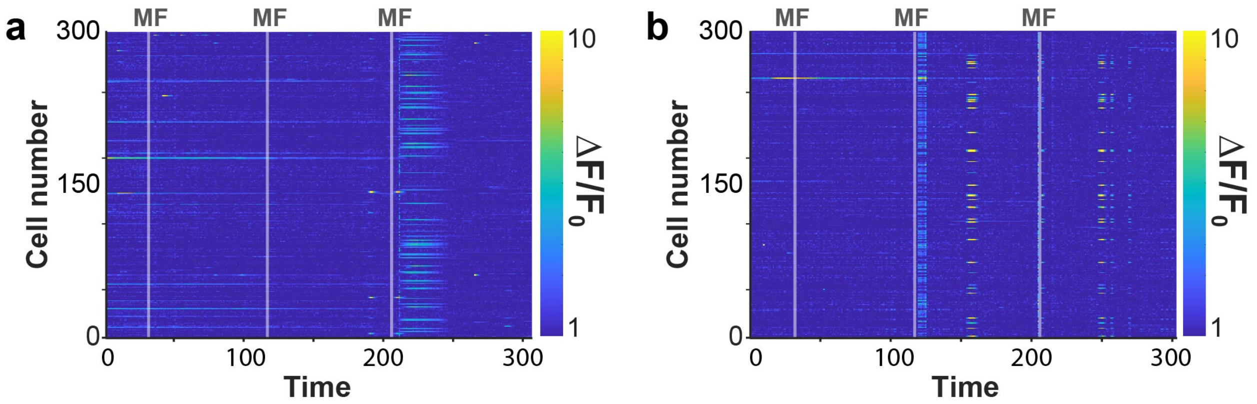


**Fig. S20| GCaMP6s imaging of neurons in the presence of drugs. a, b** GCaMP6s $\Delta F/F_0$ in 300 neurons (n=3 plates) treated with drugs corresponding to the average $\Delta F/F_0$ plots shown in **Fig. 3g,h**. **a,** Neurons treated with tetrodotoxin (TTX, 1 µM). **b,** Neurons treated with 6-cyano-7-nitroquinoxaline-2,3-dione (CNQX, 20 µM) and (2R)-amino-5-phosphonovaleric acid (AP5, 100 µM) cocktail. Vertical grey bars mark magnetic field epochs (MF, 2s, 220 mT OMF; 10 mT, 150 Hz AMF).

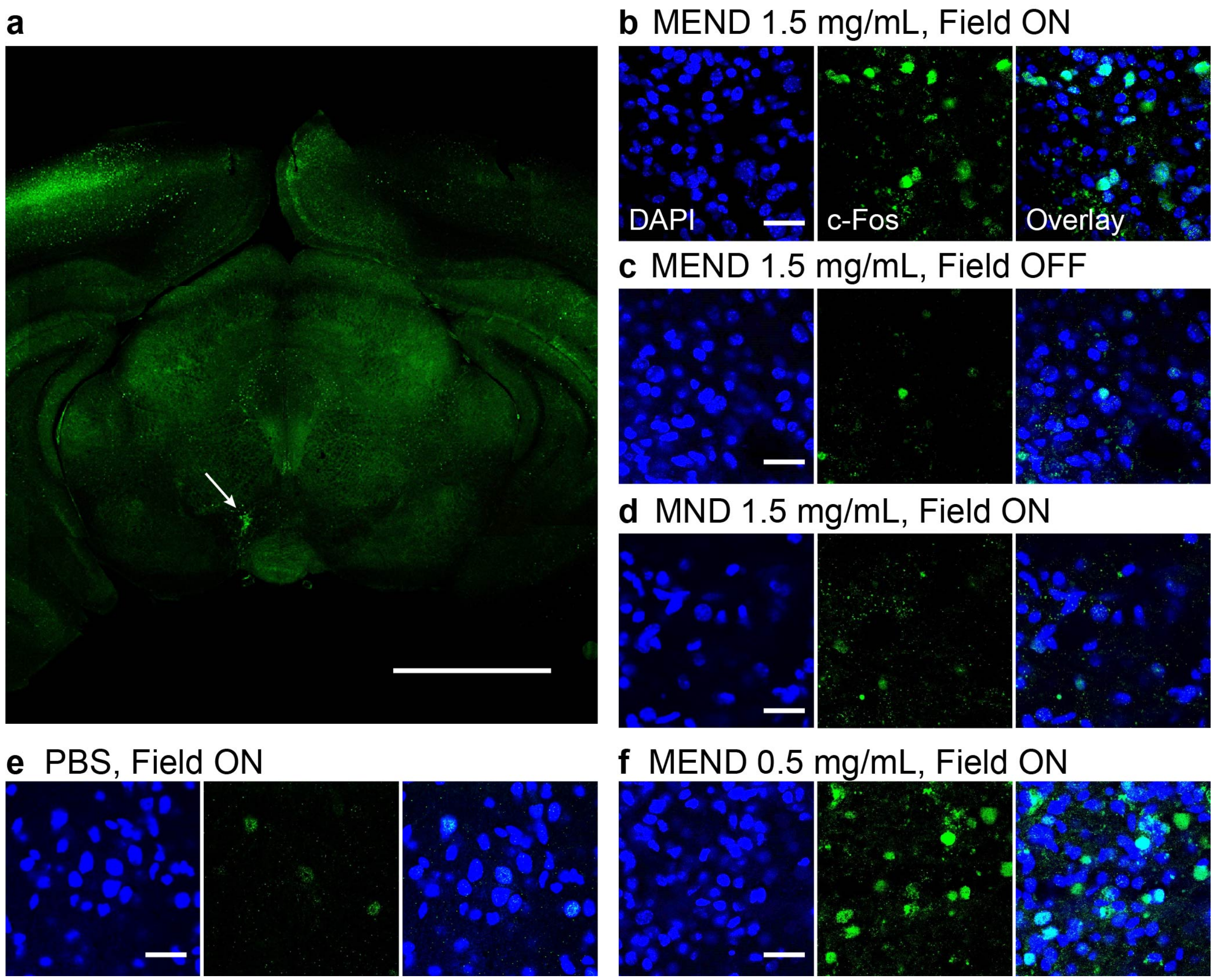


**Fig. S21| Examples of c-Fos expression in the ventral tegmental area (VTA). A,** Confocal micrograph of c-Fos expression (green) visible across the entire brain slice, with distinct increase in the VTA on the side injected with MENDs (1.5 µL at 1.5 mg/mL, marked with an arrow) following exposure to magnetic field (220 mT DC, 150 Hz, 10 mT AC; three 2s epochs separated by 90s rest epochs). Scale bar = 3 mm. **b-f** Higher resolution (20X) confocal images of nuclear stain DAPI (blue), c-Fos (green), and their overlay in brain slices from mice in different experimental groups exposed to various conditions. Scale bars = 25 µm. **b,** A mouse injected with MENDs and exposed to magnetic field. **c,** A mouse injected with MENDs and not exposed to magnetic field. **d,** A mouse injected with control $Fe_3O_4$ nanodiscs (MNDs, 1.5 µL at 1.5 mg/mL) and exposed to magnetic field. **e,** A mouse injected with phosphate buffered saline (PBS, 1.5 µL) and exposed to magnetic field. **f,** A mouse injected with a low concentration of MENDs (1.5 µL at 0.5 mg/mL) and exposed to magnetic field.

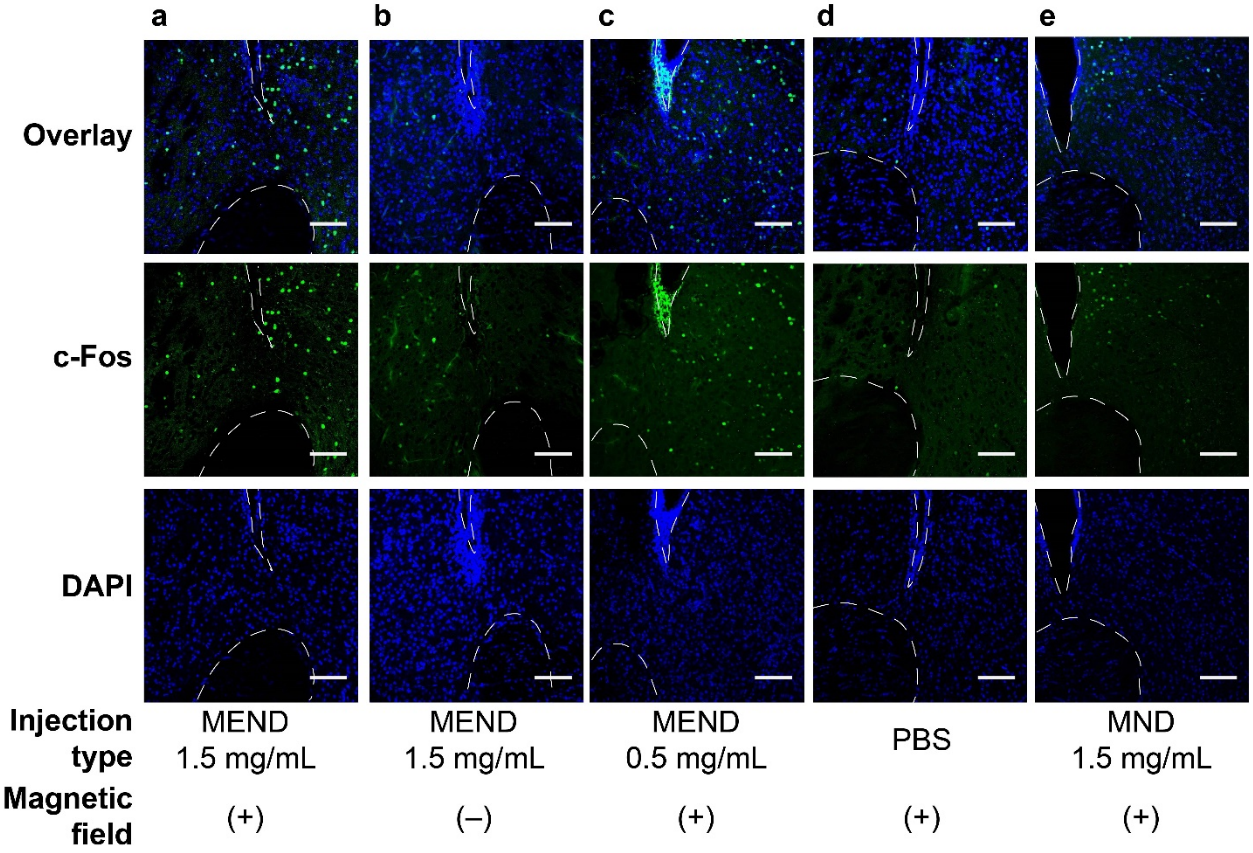


**Fig. S22| Confocal images of c-Fos expression in the nucleus accumbens (NAc) at different nanomaterial injection and magnetic field application conditions. a,** A mouse injected with MENDs (1.0 mg/mL, 1.5 µL) and exposed to magnetic field (220 mT OMF; 150 Hz, 10 mT AMF, three 2 s epochs separated by 90 s rest epochs). **b,** A mouse injected with MENDs and not exposed to magnetic field. **c,** A mouse injected with a low concentration of MENDs (0.5 mg/mL, 1.5 µL) and exposed to magnetic field. **d,** A mouse injected with PBS (PBS, 1.5 µL) and exposed to magnetic field. **e,** A mouse injected with control MNDs (1.0 mg/mL, 1.5 µL) and exposed to magnetic field. c-Fos is marked with green, DAPI is marked with blue. Scale bars = 100 µm.

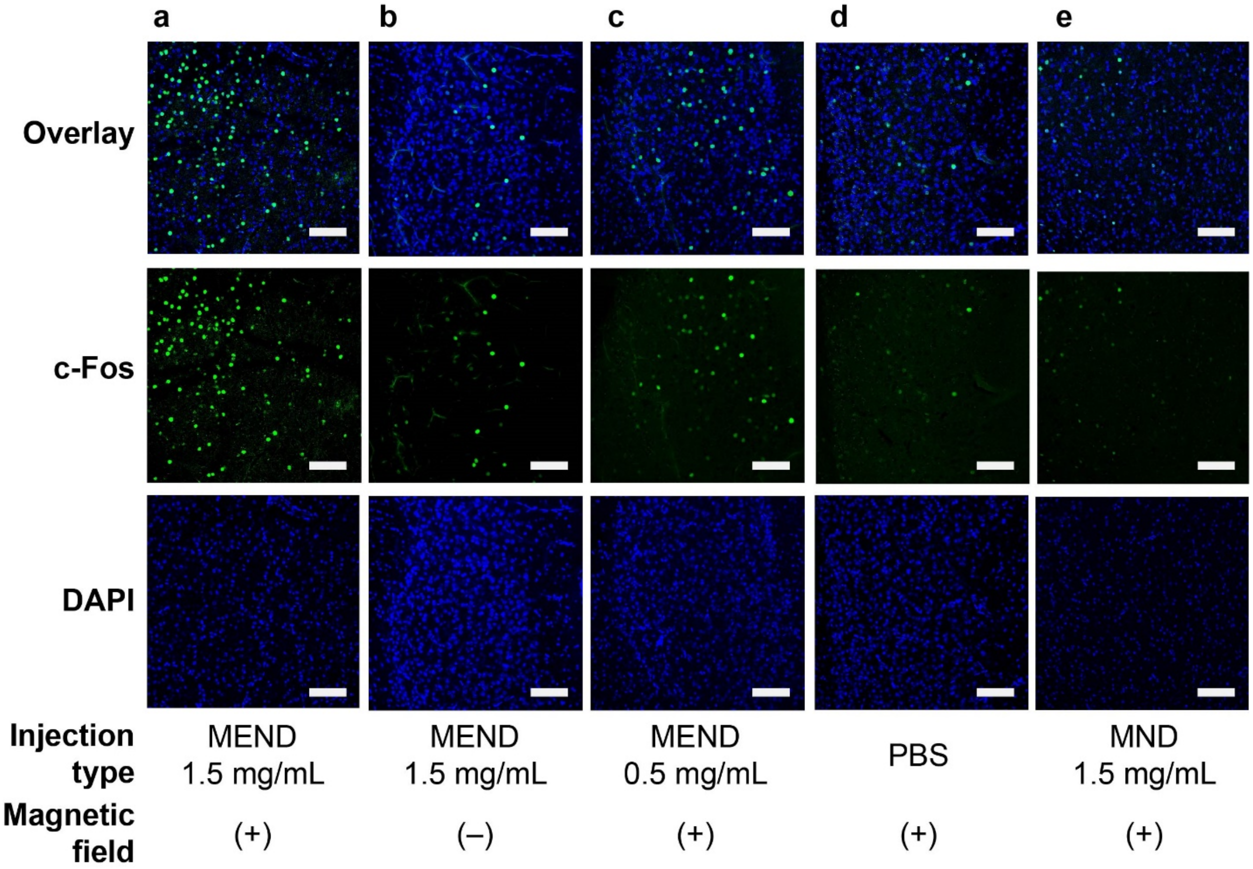


**Fig. S23| Confocal images of c-Fos expression in the medial prefrontal cortex (mPFC) at different nanomaterial injection and magnetic field application conditions. a,** A mouse injected with MENDs (1.0 mg/mL, 1.5 μL) and exposed to magnetic field (220 mT OMF; 150 Hz, 10 mT AMF, three 2 s epochs separated by 90 s rest epochs). **b,** A mouse injected with MENDs and not exposed to magnetic field. **c,** A mouse injected with a low concentration of MENDs (0.5 mg/mL, 1.5 μL) and exposed to magnetic field. **d,** A mouse injected with PBS (PBS, 1.5 μL) and exposed to magnetic field. **e,** A mouse injected with control MNDs (1.0 mg/mL, 1.5 μL) and exposed to magnetic field. c-Fos is marked with green, DAPI is marked with blue. Scale bars = 100 μm.

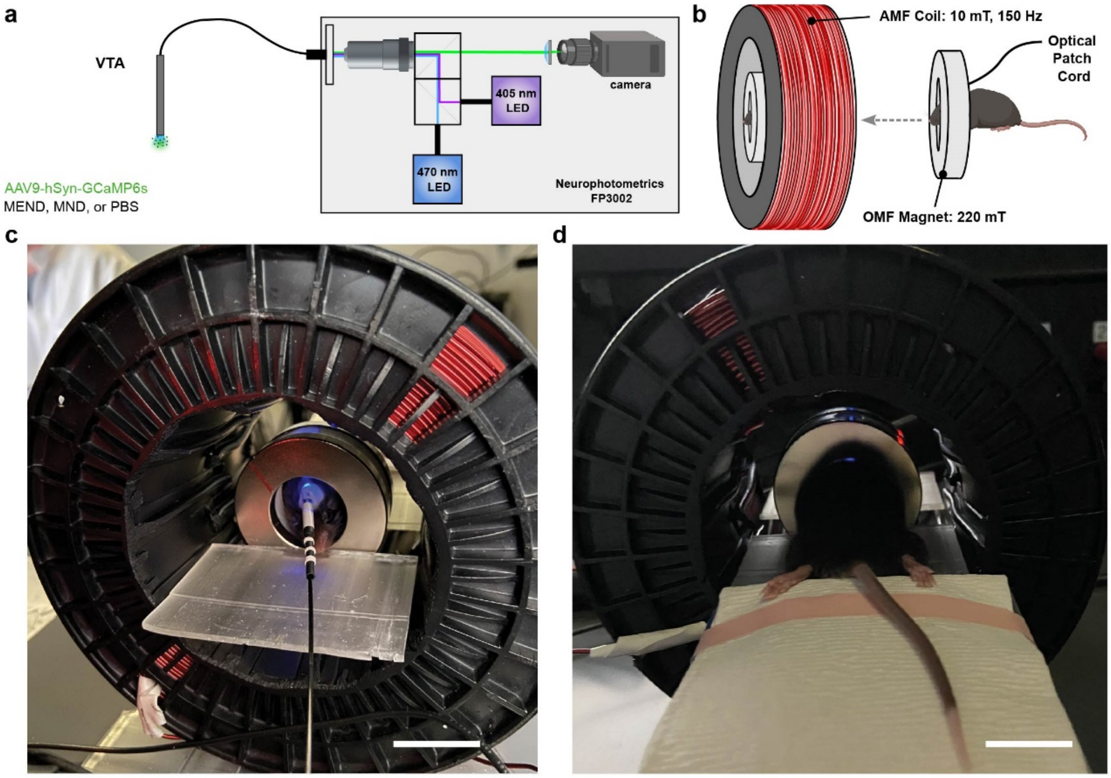


**Fig. S24| Assay for fiber photometry combined with MEND coil setup. a,** An illustration of the fiber photometry system. **b,** An illustration of magnetic field generation system. **c, d,** Photographs of front and back views of the experimental setup used for fiber photometry recordings during magnetoelectric stimulation (220 mT OMF; 150 Hz, 10 mT AMF) mediated with MENDs and the corresponding control experiments. Scale bars are 2 cm.

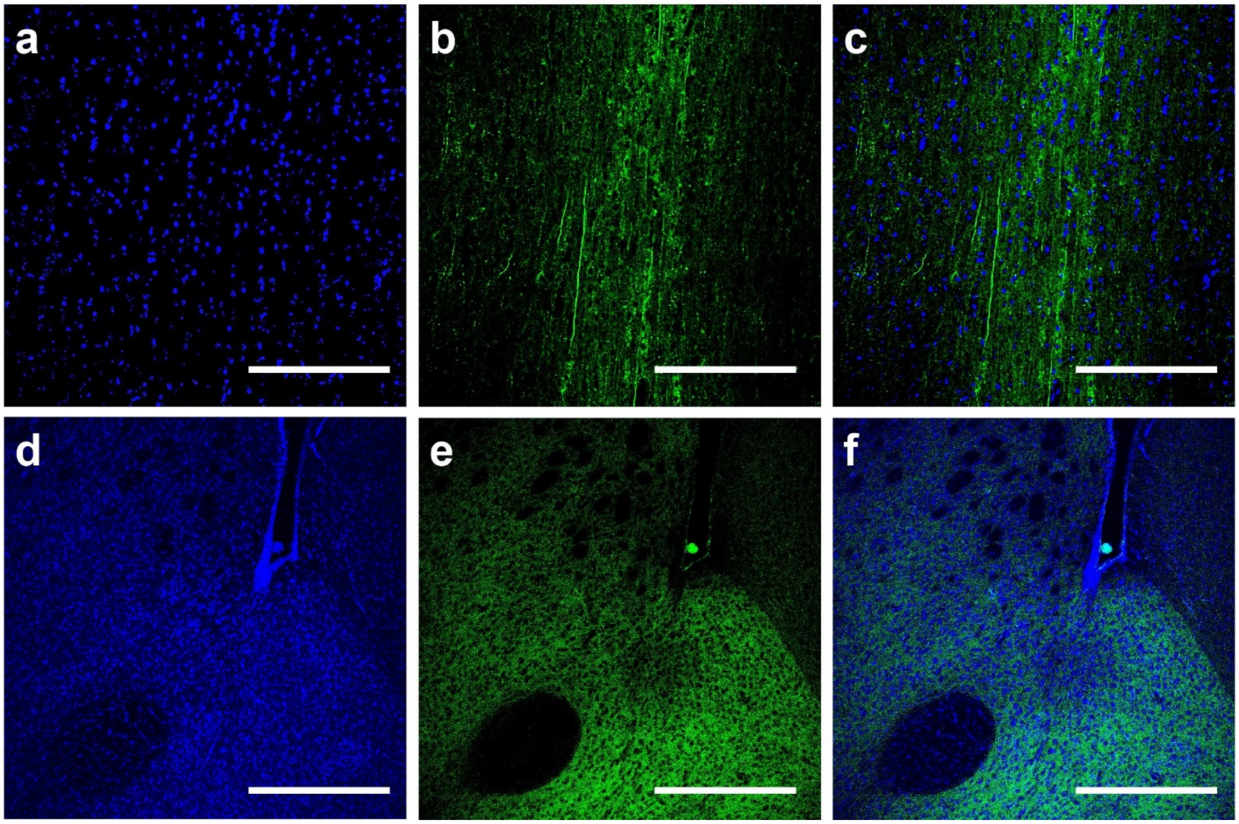


**Fig. S25| Expression of GCaMP6s on the dopaminergic circuit. a-f** Confocal images of GCaMP6s expression (green) and DAPI nuclear marker (blue) of the axons VTA projecting to mPFC (**a-c**) and NAc (**d-f**). **a, d**, DAPI-stained cells. **b, e,** show the GCaMP6s-expressing axons. **c, f,** Overlay of DAPI and GCaMP6s signals. Scale bars = 150 μm.

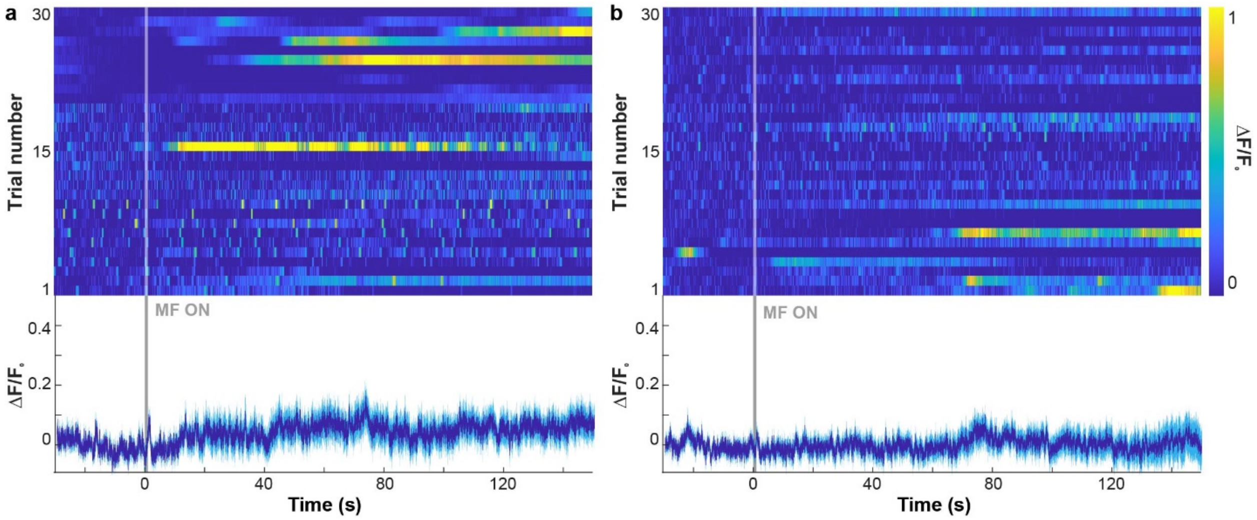


**Fig. S26| Fiber photometry of GCaMP6s fluorescence in vivo. a, b,** Photometric recordings of average relative GCaMP6s fluorescence in the VTA of the mice injected with **a,** MND (1.5 mg/mL) and **b,** PBS. Dark blue lines represent average and cyan shading represents standard deviation (n=3 mice per condition; 10 trials per mouse). Vertical grey bars denote magnetic field epochs (2 s, 220 mT OMF; 150 Hz, 10 mT AMF).

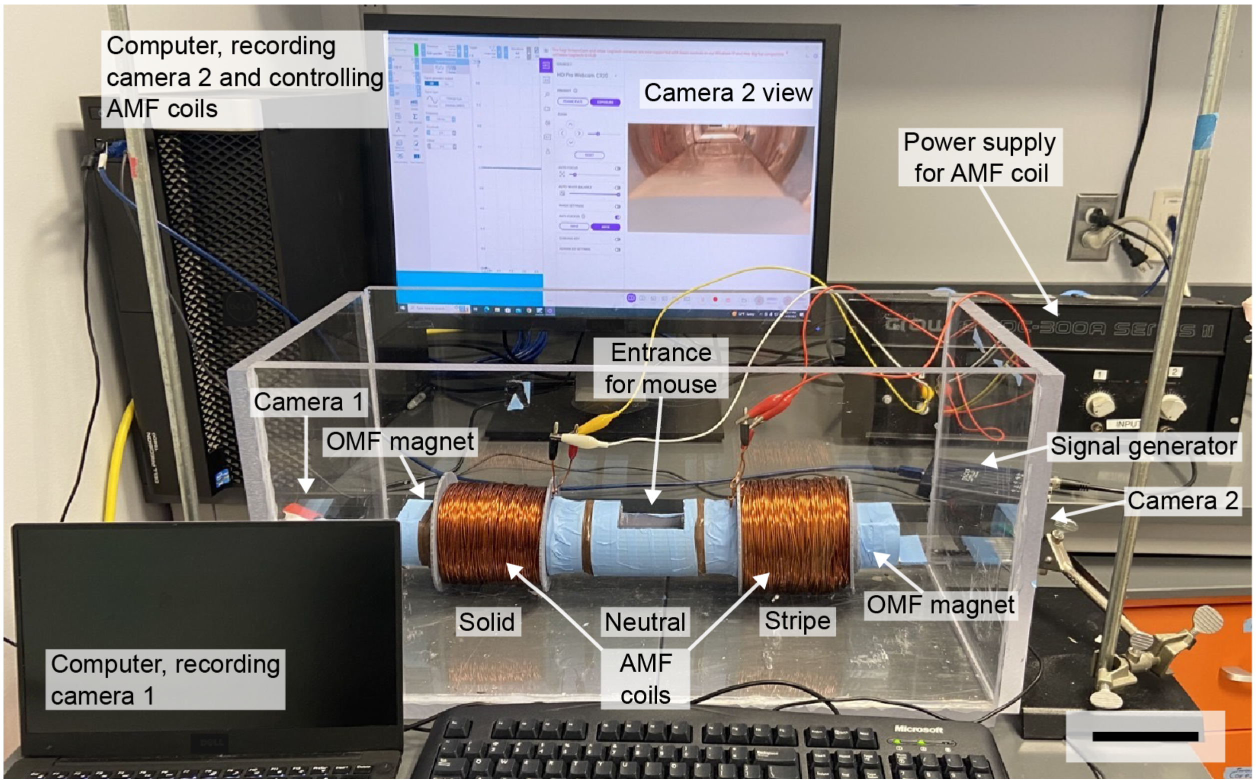


**Fig. S27| Setup for the behavior assays.** A photograph of the custom apparatus employed for place-preference assays enabling application of magnetic fields necessary for magnetoelectric neuromodulation. Scale bar is 10 cm.

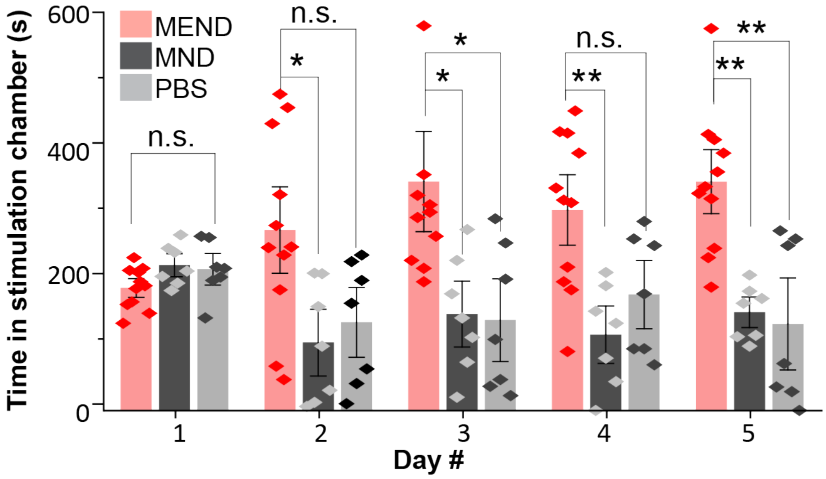


**Fig. S28| Summary of place-preference assays.** Time spent by each subject in the stimulation chamber of the arena, out of a total test time of 600 s for each trial. Scale bars denote standard deviation. (n=11 for MEND group, n=7 for MND and PBS, Kruskal-Wallis test and Tukey's post-hoc comparison test, ***$P < 0.001$, **$P < 0.01$, *$P < 0.05$, n.s. $P>0.05$). The concentration of MENDs and MNDs was 1.5 mg/mL, all injections were 1.5 µL.

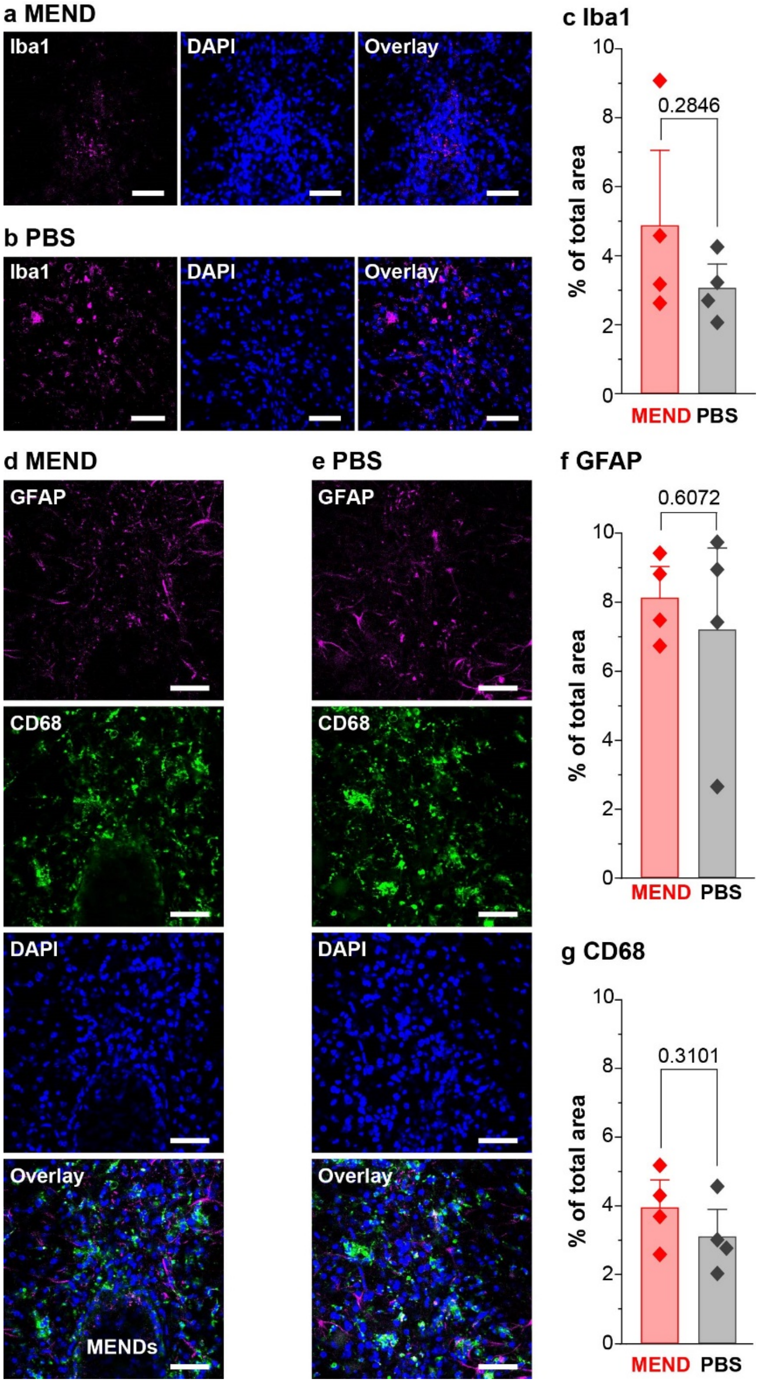


**Fig. S29| Biocompatibility assessment of MENDs (1.5 µL at 1.5 mg/mL) unilaterally injected into the VTA as compared to PBS injections contralaterally. a, b, d, e,** Representative confocal images and **c, f, g,** average percentages of expression of **a-c,** Iba1, **d,f,** GFAP (**d,f**), and **e, g,** CD68 in the brains of mice 2 weeks following surgery for unilateral injection with MENDs and contralateral injection with PBS. 90 min before the perfusion, the animals were applied to the magnetic field (220 mT DC and 150 Hz, 10 mT AC; three 2 s epochs separated by 90 s rest epochs). Scale bars = 50 µm. Two-sample t-test was performed (n=4 mice per group, P values are noted on plots in panels **c, f,** and **g**).

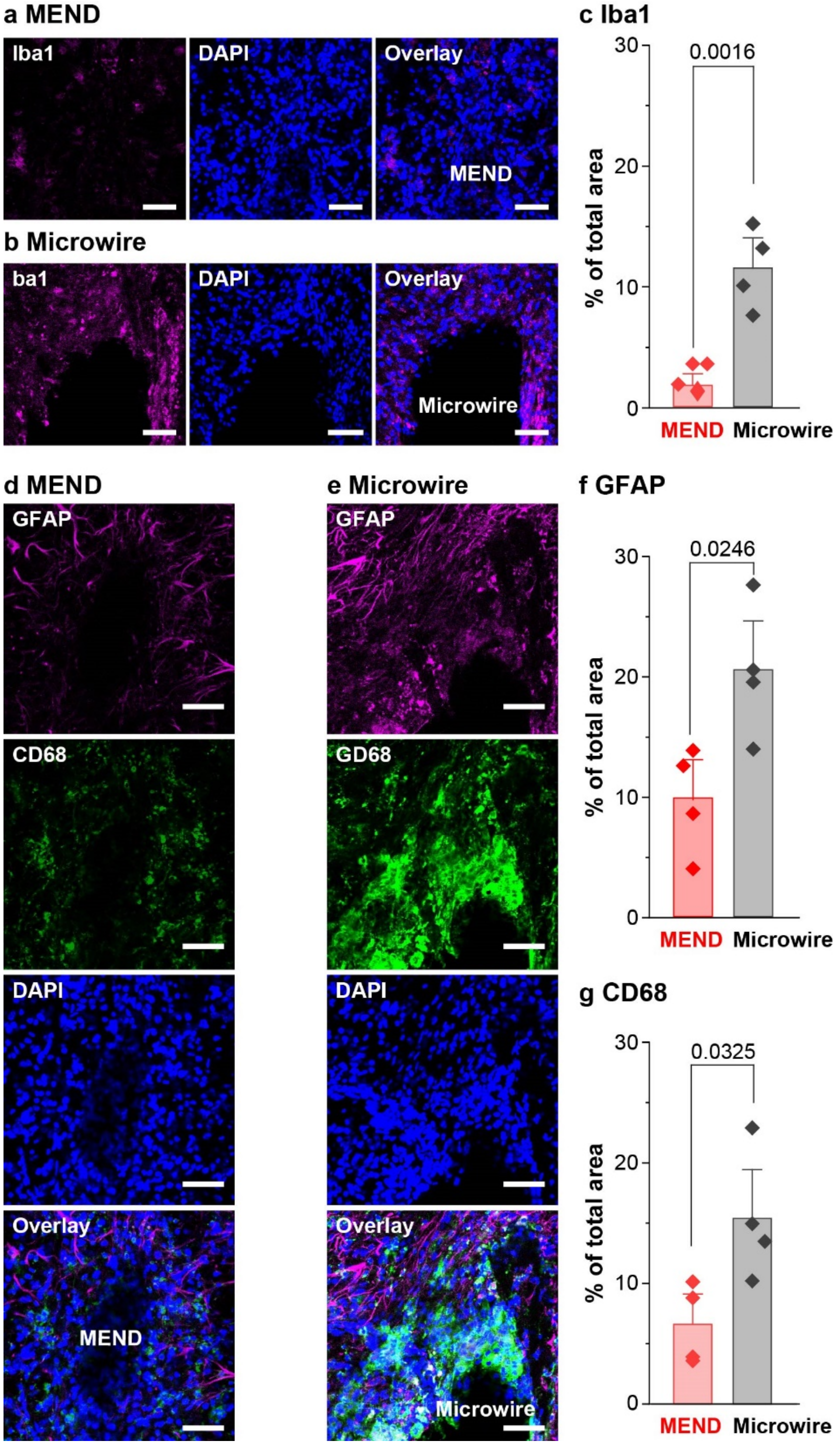


**Fig. S30| Biocompatibility assessment of MENDs (1.5 µL at 1.5 mg/mL) unilaterally injected into the VTA as compared to a 300 µm stainless steel microwire implanted contralaterally.** Representative confocal images (**a, d, e**) and average percentages (**c, f, g**) of expression across conditions of Iba1 (**a-c**), GFAP (**d, f**), and CD68 (**e, g**) in the brain of mice 2 weeks following surgery for unilateral MENDs injection and contralateral implantation of 300 µm stainless steel microwire. Scale bars = 50 µm. Two-sample t-test was performed (n=4 mice per group, P values are noted on plots in panels c, f, and g).

**Supporting Videos**

**Supporting Video 1.** GCaMP6s fluorescence changes (10X speed) in primary hippocampal neurons decorated with MENDs in response to 10s pulses of combined offset magnetic field (OMF) 220 mT and alternating magnetic field (AMF) 10 mT, 150 Hz.

**Supporting Videos 2 and 3.** GCaMP6s fluorescence changes (real time) in primary hippocampal neurons decorated with MENDs in response to 10s magnetic field epochs with the variation in the AMF frequency, while maintaining its amplitude (10 mT) and the magnitude of OMF (220 mT).

**Supporting Video 4.** GCaMP6s fluorescence changes (real time) in primary hippocampal neurons decorated with MENDs in response to 2 s epochs of combined 220 mT OMF and 10 mT, 150 Hz AMF. Separation between stimulation epochs was 120s, 90s, 60s, 30s, and 10s, and each stimulation sequence was repeated three times.

**Supporting Video 5.** GCaMP6s fluorescence changes (real time) in primary hippocampal neurons decorated with MENDs in response to 2 s epochs of 220 mT OMF and 10mT, 150 Hz AMF in the presence of 1 μM tetrodotoxin.

**Supporting Video 6.** GCaMP6s fluorescence changes (real time) in primary hippocampal neurons decorated with MENDs in response to 2 s epochs of 220 mT OMF and 10mT, 150 Hz AMF in the presence of 20 μM 6-cyano-7-nitroquinoxaline-2,3-dione (CNQX) and 100 μM (2R)-amino-5-phosphonovaleric acid (AP5).
